# Rapid and dynamic evolution of a giant Y chromosome in *Silene latifolia*

**DOI:** 10.1101/2023.09.21.558759

**Authors:** Takashi Akagi, Naoko Fujita, Kanae Masuda, Kenta Shirasawa, Kiyotaka Nagaki, Ayano Horiuchi, Eriko Kuwada, Riko Kunou, Koki Nakamura, Yoko Ikeda, Koichiro Ushijima, Deborah Charlesworth

**Affiliations:** Graduate School of Environmental and Life Science, Okayama University, Okayama, Japan; Japan Science and Technology Agency (JST), PRESTO, Kawaguchi-shi, Saitama 332-0012, Japan; Kazusa DNA Research Institute, Kazusa-Kamatari, Kisarazu, Chiba, 292-0818, Japan; Institute of Plant Science and Resources, Okayama University, Kurashiki, Okayama, Japan; Institute of Ecology and Evolution, University of Edinburgh, Charlotte Auerbach Road, Edinburgh EH9 2FL, United Kingdom

## Abstract

To test hypotheses about the evolution of massive sex-linked regions in plants, we sequenced the genome of *Silene latifolia*, whose giant heteromorphic sex chromosomes were first discovered in 1923. It has long been known that the Y consists mainly of a male-specific region which does not recombine with the X in male meiosis, and that this region carries the primary sex-determining genes, and other genes contributing to male functioning. However, only with a whole Y chromosome assembly can the candidates be validated experimentally, as we describe. Our new results also illuminate the genomic changes as the ancestral chromosome evolved into the current XY pair, testing ideas about why large regions of sex-linkage evolve, and the mechanisms creating the present recombination pattern.

**One sentence summary:** Based on the whole genome sequences of *Silene latifolia*, a model species for plant sex chromosome evolution, we describe discovery of genes underlying male-female flower differences, and relate the results to ideas about the evolution of the vast non-recombining regions of the Y chromosome.

## Main Text

Chromosomal sex determination is common not only in animals but also in a diversity of plant species. In contrast to many animal taxa, most extant flowering plants (or angiosperms) are functional hermaphrodites. Plant sex chromosomes evolved much more recently than those of many animals, and independently in different lineages (reviewed in 1), and are therefore interesting for studying sex chromosome establishment when separate sexes evolve from ancestral hermaphroditism. Close linkage between sex-determining genes may be involved (2), and selection for sexual dimorphism, involving traits that benefit only one sex (sexually antagonistic, or SA, traits), can lead to subsequent establishment of polymorphisms closely linked to the sex-determining factor(s), favouring further recombination suppression (3). These ideas can explain extensive male-specific regions of Y chromosomes (MSYs), that do not recombine with the X chromosome, and become differentiated from the X. However, physically extensive MSYs do not require sexually antagonistic selection. For example, sex determining factors may evolve within a genome region that was already recombinationally inactive (4). It has also been suggested that inversions preventing recombination that simply spread by chance (and carry no SA factors) may be more likely to persist on sex chromosomes than autosomes (5,6). Moreover, sexual dimorphisms do not require SA factors separate from the sex determining factors themselves (7). The plant, *Silene latifolia*, is well suited for testing such ideas.

The first observation of sex chromosomes in a flowering plant was in *S. latifolia*, whose karyotype is 2n = 22A+XY (8). Its “giant” Y chromosome (approx. 500Mb, see Fig. 1a) have been a model for studying heteromorphic sex chromosomes with large non-recombining regions. After decades of cytogenetic and theoretical studies, empirical studies using limited numbers of molecular markers (9,10) revealed a pattern of Y-X sequence divergence resembling the “evolutionary strata” in humans (11), with divergence lowest for genes closest to the pseudo-autosomal region that still recombines. This indicates different episodes of Y chromosome recombination cessation. However, there are important gaps in our understanding of this Y chromosome, including why the Y-linked region is so large, and to what extent it has undergone genetic degeneration, like mammalian Y chromosomes (12). Recent advances in genome sequencing have yielded data suggesting that partial genetic degeneration has evolved (13,14), and possibly dosage compensation (15–18), but degeneration cannot be accurately quantified without an assembled Y chromosome.

**Figure 1.**
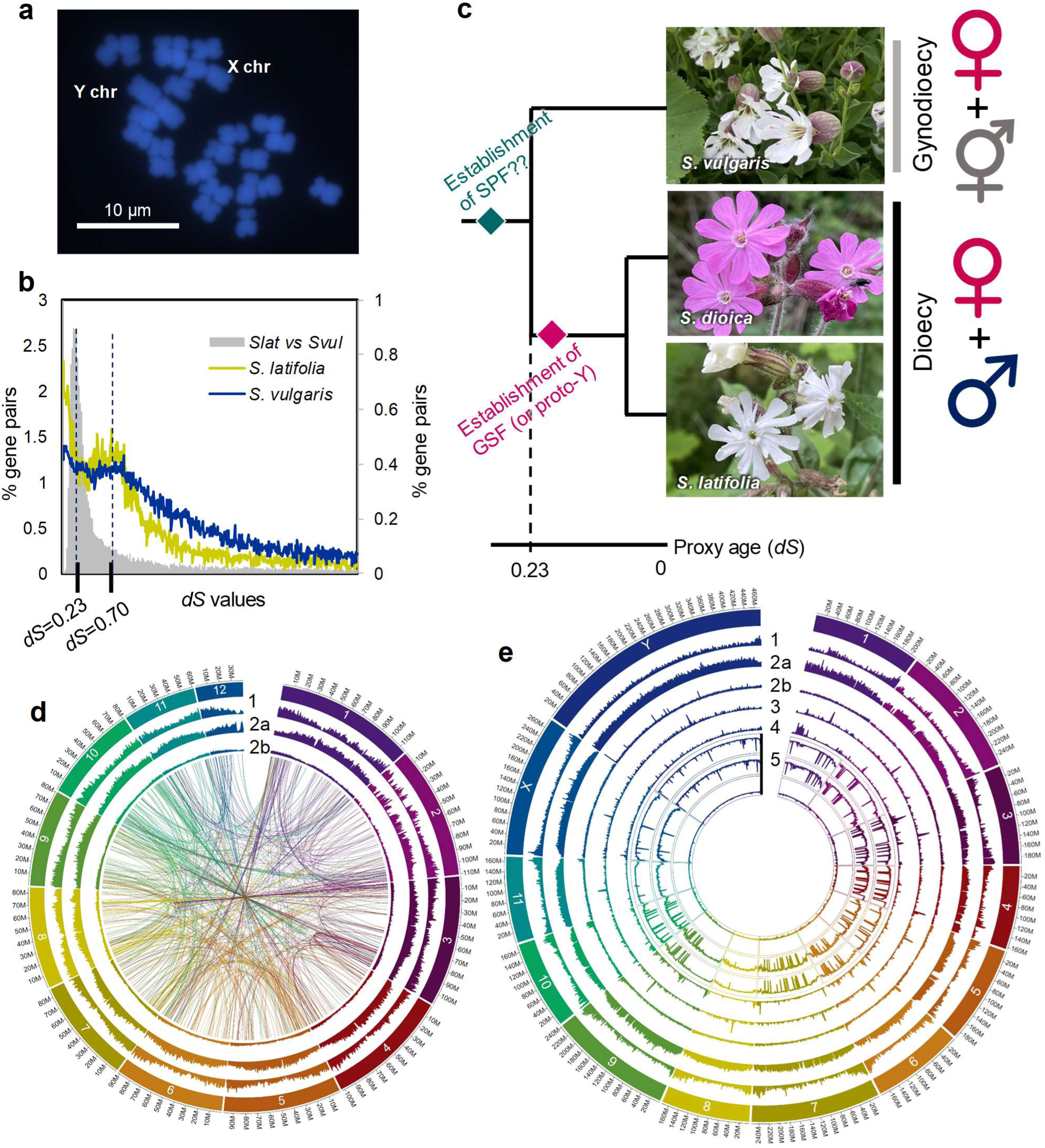
Overview of the sex chromosomes and the genome characteristics in *S. latifolia* and in an outgroup species, *S. vulgaris*. **a**, Cytogenetic observation of the giant Y and X chromosomes with DAPI staining. **b**, Estimated evolutionary context of the two sex determining factors, SPF and GSF, in dioecious species (*S. latifolia* and its close relative *S. dioica*). **c**, Histogram of the pairwise *dS* values between paralogous genes within each genome of *S. latifolia* and *S. vulgaris*, and between orthologous genes in these two species (*Slat vs Svul*). The interspecific *dS* values exhibited a peak at *dS* = 0.23, which corresponds the proxy age for divergence of *S. latifolia* and *S. vulgaris*. **d-e**, Bird-eye views of the genomes of *S. vulgaris* (**d**) and *S. latifolia* (**e**). Circle layer 1, Gene density; layer 2, Transposable elements density (2a, LTR-type TEs; 2b, non-LTR); layer 3, standardized CENH3/H3 amount; layer 4, standardized H3K9me2/H3 amount; layer 5, weighted DNA methylation rate (5a: CG; 5b, CHG; 5c, CHH). Synteny of the gene pairs with *dS* = 0.7-0.75 was given in the center area of panel **d**.

Observations of Y chromosome deletions revealed that the *S. latifolia* Y carries two sex-determining factors, a stamen-promoting factor (SPF) and a gynoecium suppressing factor (GSF) (19). This Y therefore fits the “two-mutation model” model (20), in which a loss-of-function mutation in an SPF in an ancestral hermaphrodite species creates females, and males arise by a dominant femaleness suppressing mutation, GSF, in a closely linked gene. In several dioecious plants, two genes have been detected, or inferred, and they are indeed within small genomic regions (21–24) very different from *S. latifolia*’s Y. In this model, selection favors closer linkage between the two genes involved, potentially explaining subsequent loss of Y-X recombination in the region, which is predicted to allow repetitive sequences to accumulate rapidly (25, 26), potentially forming a giant Y.

The *S. latifolia* sex chromosomes evolved after divergence from *S. vulgaris*, a gynodioecious species (with female and hermaphrodite individuals, and 2n = 24A chromosomes) (Fig. 1b). Here, we assembled chromosome-scale whole genome sequences of a *S. latifolia* male and a *S. vulgaris* hermaphrodite, using PacBio HiFi reads, Bionano optical mapping, and anchoring with cytogenetic markers, and genetic maps in both species (Fig. 1c-e; tables S1-2 detail sequence and assembly quality statistics). BUSCO (27) analysis detected 92.9% and 97.6% of universal single-copy orthologous genes in eudicotyledonous plants in *S. latifolia* and *S. vulgaris*, respectively (table S1). Synonymous site divergence between these species (*dS*) averages 0.23. Both genomes are diploid, though we detected a paleo-genome duplication event (with *dS* = 0.7, Fig. 1c), consistent with frequent plant-specific polyploidization at the K-Pg boundary (28, 29).

The *S. latifolia* Y chromosome sequence forms 6 super-contigs, totalling approimately. 480 Mb. Genetic mapping of markers in male meiosis revealed a large non-recombining male-specific region (MSY), and a pseudo-autosomal region (PAR) at one chromosome end (10,15). We therefore anchored contigs cytogenetically, by fluorescence *in situ* hybridization (FISH) using MSY region probes (Fig. 2a-h, fig. S1).

**Figure 2.**
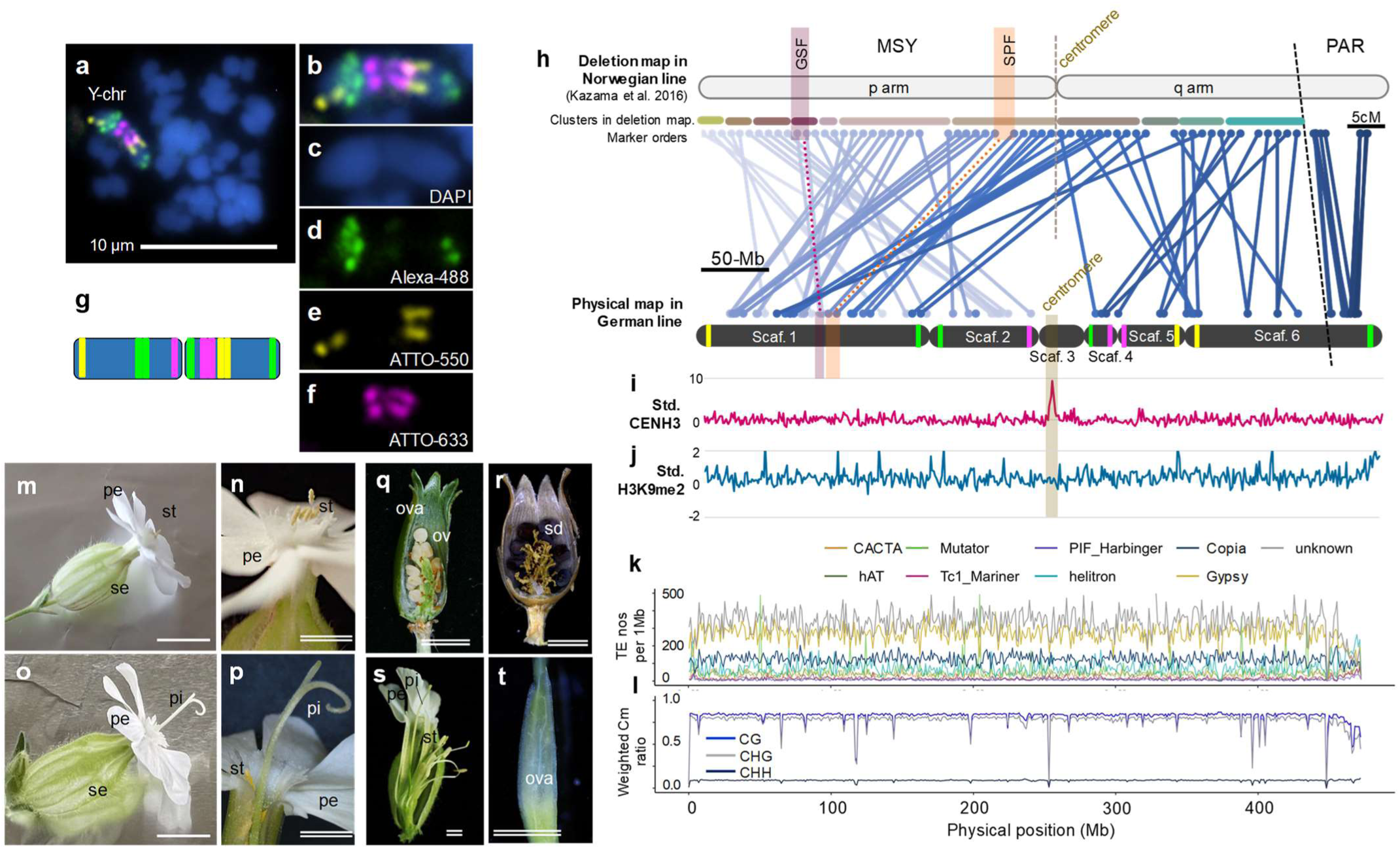
Genomic context of the *S. latifolia* Y chromosome. **a-g**, FISH anchoring of the genomic contigs. **a**, Detection of the Y chromosome by myTags probes. **b**, Enlarged image of the Y chromosome in panel **a**, merging the signals with DAPI. (**c**) Staining for three probes,Alexa-488 (**d**), ATTO-550 (**e**), and ATTO-633 (**f**). **g**, Ideogram of the Y chromosome based on the signals. **h-l**, Genomic context of the Y chromosome. **h**, Comparative analysis of the deletion map in a Norwegian line (30) (with only marker orders given, as no assembly was available) and our genome sequence of a German line. The regions including the two sex-determining genes are highlighted. In the recombinationally active PAR, the genetic map positions from Bergero et al. (10) correlate with the physical positions in our genome sequences. (**I and j**) CENH3 content (standardized by H3 amounts) (**i**) exhibits a clear single peak, whereas the standardized H3K9me2 levels (**j**) fluctuate, but show no major pattern along the chromosome. **k**, TE distribution in 1 Mb windows. LTR-type TEs, especially *Gypsy*- and *unknown*-classes, show high densities in the MSY, compared with the PAR. CG and CHG DNA methylation levels show a similar pattern (**l**). **m-t**, VIGS of *Y-SlCLV3* resulted in the production of hermaphrodite flowers in a genetically male *S. latifolia*. Control male flowers (**m-n**) exhibited functional stamens (st) but no elongated pistils. ALSV-induced gene-silencing resulted in elongation of functional pistils (pi) with normal stamens in male plants (**o-p**), producing ovules (ov) (**q**) to be viable seeds (sd) (**r**). ALSV-infected male lines often induced imperfect sterile pistils (**s-t**). pe petal, se sepal, ova ovary. Bars = 1 cm (single line), 5 mm (double line).

The Y chromosome is metacentric, but the arrangement in our family (using parent plants from Germany) differs from that in a Norwegian family (named K-line) in which marker orders were determined by deletion mapping (30) (Fig. 2h, fig. S2). The Y chromosome has clearly been rearranged within *S. latifolia*. The autosomes are also metacentric, consistent with their genetic maps, and show CENH3 antibody peaks and dense tandem repeats indicating their centromeric locations (fig. S3-S4). No matured repeats were found in the Y centromeric region (Figs. 2i and fig. S4b), which suggests frequent changes of the Y chromosome centromere location. Consistent with this, the densities of H3K9me2 (a common heterochromatin epimark) and DNA methylation in young leaves (mainly in CG and CHG contexts) are distributed evenly throughout the MSY (Fig. 2j-l). LTRs of the *Gypsy* and *unknown* classes are remarkably enriched in the non-recombining MSY and in autosomal low-recombination regions (*r* = -0.65∼-0.66, fig. S5 for all chromosome recombination rate/TEs/DNA methylation, fig. S6 for correlation test, table S3 for TE distributions), as discussed below.

Modern deletion mapping confirmed Westergaard’s finding of three MSY regions containing factors affecting male and female flower organs (31, 32). We used more precise locations identified by Kazama et al. (30), and identified regions syntenic with the GSF and SPF regions in our assembly; they included only 31 and 62 candidates for GSF and SPF, respectively (fig. S7-S8 for genomic structures of GSF and SPF regions, table S4-5 for GSF/SPF candidates). Expression analysis in early flower primordia suggested a *CLAVATA3*-like gene (*SlCLV3*) as the GSF candidate (33). Its X and Y copies have *dS*_XY_ = 0.179. Experimental virus-induced gene silencing (VIGS) of Y-encoded *SlCLV3* in male *S. latifolia* with apple-latent spherical virus (ALSV) (34) resulted in the formation of functional hermaphrodite flowers (Fig. 2m-s, fig. S9, table S6), validating this conclusion. We also identified a candidate for the SPF, an ortholog of the immunophilin-like *FKBP42/TWISTED DWARF1* (*TWD1*) gene, which is required for androecium development in other angiosperms (35,36) (fig. S10a for phylogeny). This gene is Y-specific in *S. latifolia* and its close dioecious relative, *S. dioica* (fig. S10b), and exhibits high expression in male flower primordia (table S5, fig. S8). These results are consistent with the “two-mutation” model, in which the X-linked allele of the SPF gene is non-functional.

Much of the MSY sequence shows little homology with the X (fig. S11). Gene-based syntenic block analyses also detected many Y-specific duplications (Fig. 3a-c, fig. S12); these paralogs exhibited a peak of synonymous site divergence, *dS*, at 0.241 ± 0.022 (Fig. 3b), similar to the *S. latifolia* - *S. vulgaris* value of 0.233 (Fig. 1c). This suggests that the Y chromosome stopped recombining roughly when the two lineages split, and that duplications immediately started accumulating. These duplicates have much higher non-synonymous site divergence (*dN*) than for genome-wide paralogous pairs with similar *dS* values (0.158 < *dS* < 0.316); the respective mean *dN/dS* values are 0.59 versus 0.38 (*p* = 4.6e^-26^, by a Wilcoxon’s test) (Fig. 2d). McDonald-Kreitman tests (37) indicate that most of them show nearly neutral evolution (Neutrality Index (NI) = 0.8∼1.5, *p* > 0.1 in G-tests), consistent with the predicted weaker selection after duplication (38); some genes were selected against, and a few positively selected (*p* < 0.1) (Fig. 3e).

**Figure 3.**
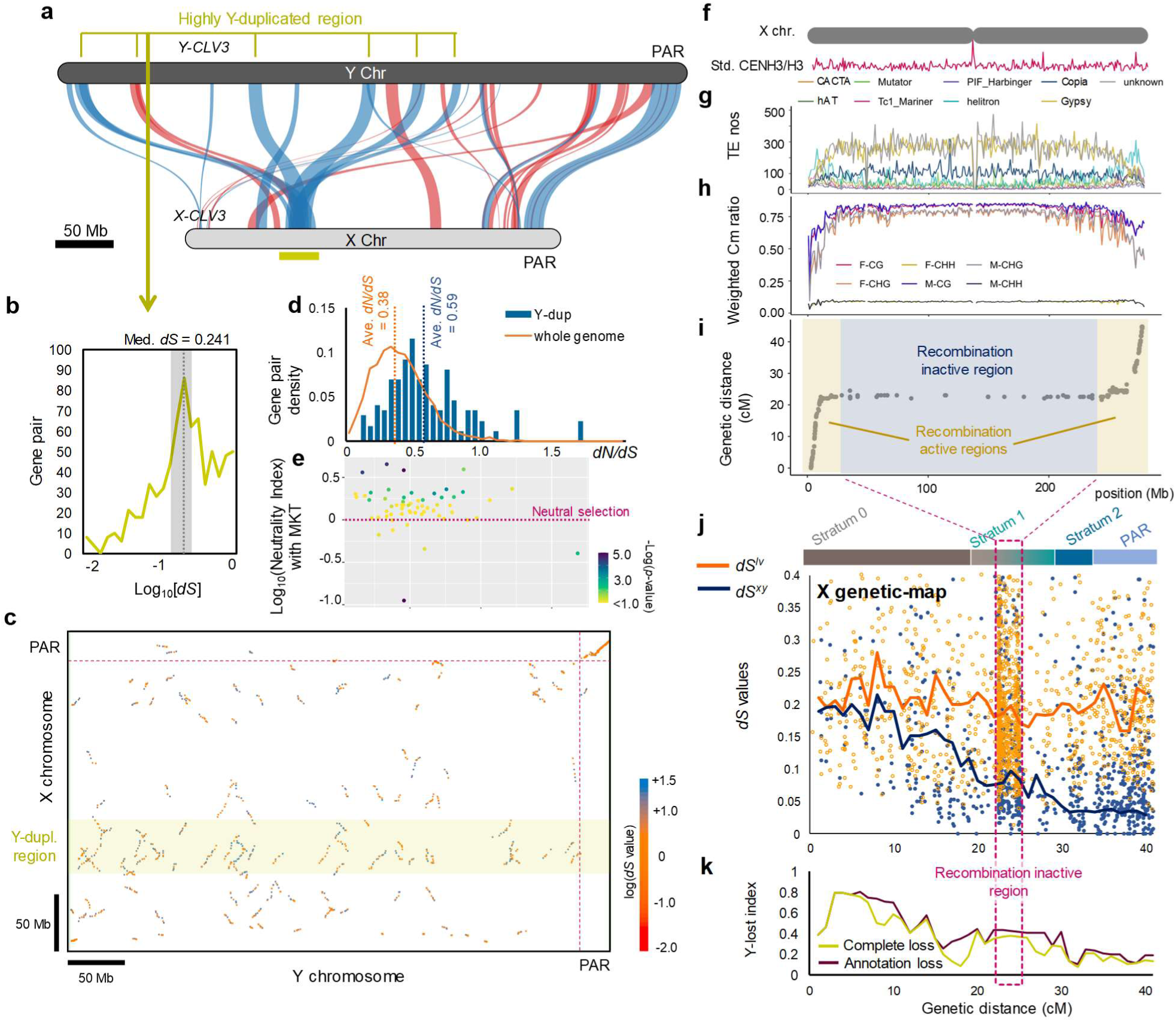
Comparative analysis of Y and X chromosomes in *S. latifolia*. **a**, Gene-order-based synteny analysis between the X and Y chromosomes. The X-linked region marked with a gold horizontal band includes sequences that are highly duplicated in the Y chromosome. **b**, Pairwise *dS* values of genes in the Y-duplicated regions. The peak with *dS* values of 0.158 to 0.316 is highlighted in gray. **c**, Locations of syntenic blocks in the X and Y chromosomes, with colours indicating the mean *dS* values between the X- and Y-linked sequences (see the key). **d**, Values of *dN/dS* in the paralogous gene pairs with *dS* in the interval between 0.158 and 0.316; pairs are shown separately for the Y-duplicated region and the whole genome. **e**, Distribution of the Neutrality Index (NI) in McDonald-Kreitman tests (the x axis values are the *dN*/*dS* values given in panel **d**). **f-i**, Characteristics of the X chromosome. **f**, standardized CENH3 amount (corrected by H3 amount). **g**, TE distribution with 1-Mb windows. **h**, Weighted DNA methylation levels. **i**, Genetic recombination map estimated using an F1 population of 228 individuals, showing that the regions with highly elevated LTR-type TEs, especially *Gypsy*- and *unknown*-classes, and CG and CHG DNA methylation levels correspoind with the recombination arrested region. **j-k**, Evolutionary strata in terms of Y-X synonymous site divergence, and of genetic degeneration. The x axis shows positions in X chromosome genetic map (fig. S13 shows the same data with positions in the X assembly). **j**, *dS* values between X- and Y-linked alleles (*dS^XY^*) and between *S. latifolia* X-linked alleles and their orthologues in *S. vulgaris* (*dS^lv^*). The lines indicate median *dS* values in 1 centiMorgan bins. Based on Pettitt’s change-point test using the estimated *dS^XY^* values, we defined three strata, 0 (*dS^XY^* > approx. 0.10), 1 (0.05 to 0.10) and 2 (0.03 to 0.05). **k**, Proportion of genes ancestrally present (defined in the text) that have been lost from the *S. latifolia* Y. Complete loss: the genome region corresponding to the gene is absent from the Y, Annotation loss: the gene is not annotated in Y.

In genetic maps, recombination in *S. latifolia* female meiosis is also infrequent across much of the X chromosome (Fig. 3i), similar to the autosomes and *S. vulgaris* chromosome maps, consistent with a recent report (39). Recombinationally inactive regions occupy most of the middle region of each chromosome, and indicate the presence of physically large pericentromeric regions previously inferred in *S. latifolia* (10, 15). Consistent with rare recombination, LTR-TE and DNA methylation densities are high in these regions, and similar to the values in the MSY (Fig. 3f-i, figs. S5-6).

To test for evolutionary strata on the sex chromosome pair, we estimated synonymous site divergence between complete X chromosome genes’ coding sequences and those of their Y-linked alleles (*dS^XY^*, Fig. 3j). Genes distant from the PAR have the highest values, similar to *dS* between *S. latifolia* and *S. vulgaris*. The values decline with proximity to the PAR, using either the X genetic or physical map positions (Fig. 3j, Fig. S13), confirming previous results that (without an X assembly), used X genetic map positions (40,41,15). Based on change-point tests (Pettitt 1979), we detected two changes in *dS^XY^* values (with *p* = 2.6e^-11^ and 2.7e^3^), defining three strata, 0-2 (Fig. 3j). Within the oldest stratum, 0, median *dS^XY^* values are 0.10∼0.23, close to the divergence between *S. latifolia* and *S. vulgaris* (*dS^lv^*). Stratum 2, adjoining the PAR, corresponds to the stratum that evolved since the split from *S. dioica* (41,42).

With the availability of a Y chromosome assembly, we can now ask whether strata are also evident in terms of gene losses from the Y. We defined ancestral genes as those present on the *S. latifolia* X and on the *S. vulgaris* homolog, following the approach developed for humans (43). The proportion of genes shared with the syntenic region of *S. vulgaris* is similar across the entire X chromosome (Fig. 3k, fig. S13). Stratum 0 corresponds to part of the X chromosome recombining region to the left of the central rarely recombining region, and includes many windows in which > 70% of *S. latifolia* genes are X-specific (Fig. 3k), revealing much more degeneration than previous estimates without a Y assembly (44). Stratum 1 starts within the same recombining X region, and spans the >200 Mb pericentromeric rarely recombining region (see Fig. 3i; genes within this small genetic map interval are within the dotted magenta box) and extends into the right-hand X recombining region. This stratum displays fewer losses of ancestral genes than Stratum 0, and they are even rarer in Stratum 2.

Chromosome-wide synteny analysis between the *S. latifolia* sex chromosomes and *S. vulgaris* confirmed the previous suggestion that the chromosomes have been rearranged in either or both species, perhaps by fusions of *S. vulgaris* chromosomes during formation of the *S. latifolia* X (10, 45). Portions of four different *S. vulgaris* chromosomes correspond to the *S. latifolia* X, and the Y chromosome is further greatly rearranged (Fig. 4a-d). Most *S. latifolia* X/Y regions are recombinationally active in *S. vulgaris* (Fig. 4d), but in *S. latifolia* most are not: a recombining part of *S. vulgaris* chromosome 4 corresponds with the region occupied by Stratum 0 and most of Stratum 1, which do not recombine with the X in males, and, in females, only part is recombinationally active, but most is within the large rarely recombining pericentromeric region.

**Figure 4.**
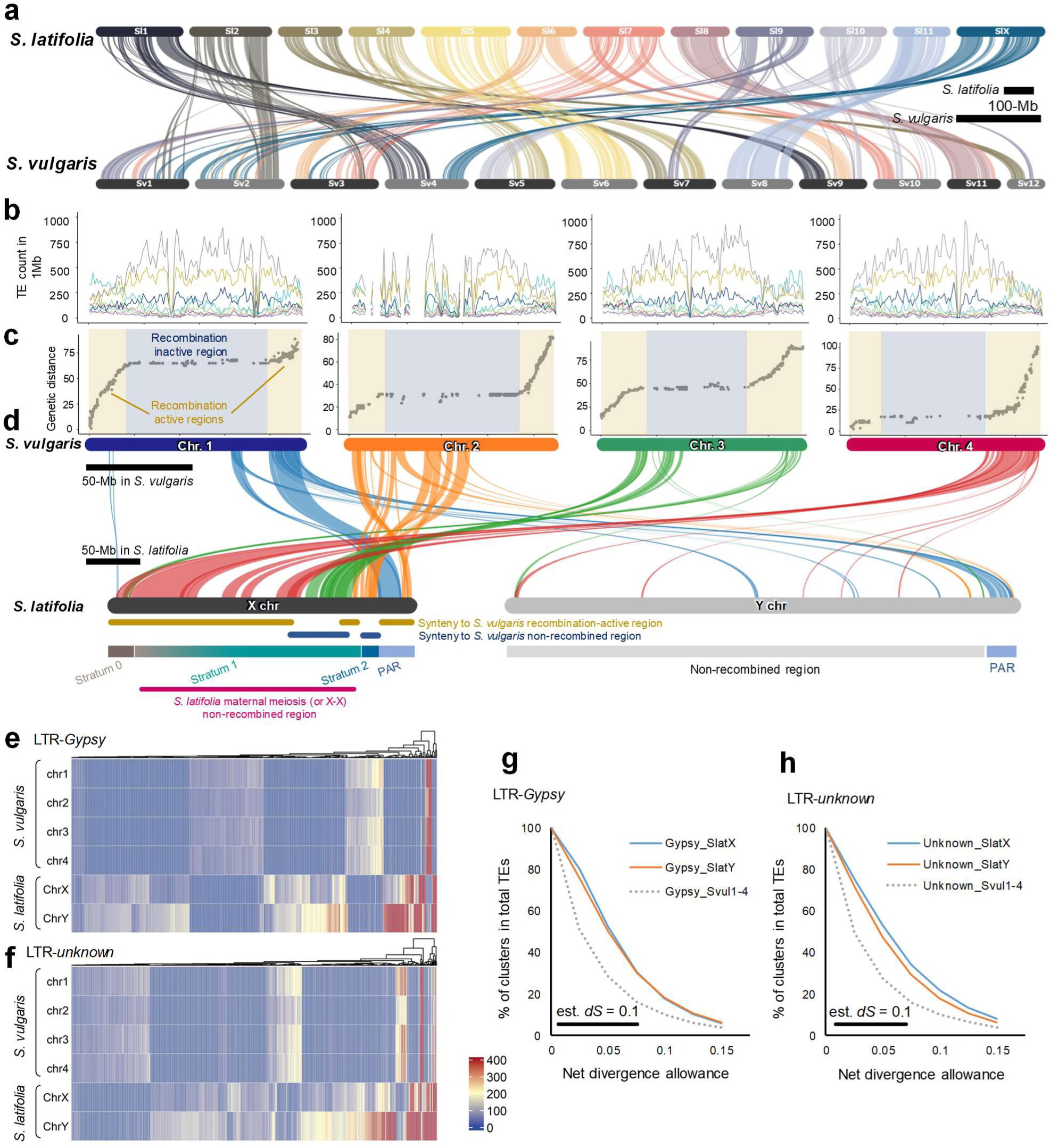
Comparative genomic analysis between *S. latifolia* and *S. vulgaris*. **a**, Whole genome synteny between *S. vulgaris* and *S. latifolia*. The sex chromosome(s) of *S. latifolia* was syntenic to *S. vulgaris* chromosome. **b-d**, Genomic contexts of the chromosome 1-4 of *S. vulgaris* and their detailed relationship to the X and Y chromosome of *S. latifolia*. **b**, TEs distribution. Refer to Fig. 3f for their annotations. **c**, Genetic recombination map. Consistent with X- and Y chromosomes, their wide recombination arrested regions were enriched with *Gypsy*- and *unknown*-classes LTR-TEs. **d**, Half of the X chromosome (and potentially also ancestral Y chromosome) was syntenic to recombination-active regions of the Chr. 4 in *S. vulgaris*. The other half was thought to be derived from chromosome fusions of Chr. 1-3 in *S. vulgaris*, of which some are currently non-recombining. **e-f**, Sequence homology-based clustering of the *Gypsy*- (**e**) and *unknown*-class (**f**) LTR-TEs extracted from X/Y chromosomes in *S. latifolia* and chromosome 1-4 in *S. vulgaris*. A large part of the clusters tended to be constituted in X- and Y-specific and lineage-specific manners. **g-h**, Numbers of *Gypsy* (**g**) and *unknown* type (**h**) TE clusters (defined as sequences with the divergence values indicated on the x axis) as proportions of the total numbers of each such TE type. The x axis shows threshold values of up to 15% for defining clusters, and these thresholds approximately quantify the divergence of extant insertions from their ancestral sequences, and therefore reflect the insertions’ ages). The figure shows that the *S. latifolia* X and Y are rich in TE bursts that are older than bursts on the *S. vulgaris* chromosome 4.

Recombination rates and TE densities are highly correlated in both *S. latifolia* and *S. vulgaris*. Recombinationally inactive regions are highly enriched with LTR-*Gypsy* and LTR-*unknown* class TEs (Fig. 4b-c, *r* = -0.68∼-0.69, fig. S5-6, fig. S14-15). To test the potential involvement of TEs in sex chromosome expansion, *Gypsy*- and *unknown*-class TEs were extracted from the *S. latifolia* X- and Y chromosome sequences and the 4 syntenic chromosome regions in *S. vulgaris*. Most sequences clustered by the species, or were specific to the *S. latifolia* X or Y (Fig. 4e-f, table S6, based on a net divergence threshold of 0.20). Even in the clusters shared by the *S. latifolia* X and Y, or by both species, recently generated sub-clusters (*dS* < 0.1) tended to be more specific to one chromosome than older ones, as expected (fig. S16, table S6). Divergence estimates of LTR coding sequences (Fig. 4g), suggest that most TE insertions occurred recently, consistent with insertions generally being deleterious (25). The insertions in the *S. latifolia* X and Y tend to be older than those in *S. vulgaris*, whereas the age estimates indicate that most TE bursts (*dS* < 0.1) occurred after the oldest stratum 0 formed (*dS* > 0.1). These TEs were therefore not the cause of recombination arrest, but probably contributed to expansion of the MSY after recombination stopped. TE accumulation may also have contributed to the large sizes of the *S. latifolia* and *S. vulgaris* pericentromeric regions.

We next consider what might have caused the suppressed recombination in males in regions that recombine between the X chromosomes in females. Under the two-gene hypothesis described above, both primary sex determining genes should be in the region that evolved suppressed recombination first. The GSF candidate is indeed within Stratum 0 (between 84-86 Mb of our Y assembly, and the terminal location of its X-linked copy probably reflects its ancestral position; Fig. 3a, fig S7). However, the SPF candidate’s ancestral position is unclear, as it has no X copy; its Y copy is nearby in our Y assembly (between 90-97 Mb, fig. S8), but Y not in the deletion map (Fig. 2h). We also found a possible sexually antagonistic gene, *SlBAM1*, near the end of Stratum 0 (fig. S17a-c), potentially contributing to *S. latifolia* secondary sexual trait dimorphism(s). *BARELY ANY MERISTEM 1/2* (*BAM1/2*)-like receptor-like kinases often function in reproductive processes and flower morphological development via *CLV1/3*-related pathways in plants (46–48). *SlBAM1* shows sex linkage (with a male-specific *Y-BAM1* sequence) in a sample of *S. latifolia* and *S. dioica* individuals, and the gene is also present in the hermaphrodite *S. conica* (fig. S17b). Ectopic expression of *Y- BAM1* in *Nicotiana tabacum*, under the regulation of its native promoter, resulted in substantially increased flower numbers per inflorescence (*p* = 0.39e^-4^) and reduced flower size (*p* = 0.00015), consistent with *S. latifolia* male-specific traits (fig. S18, table S7) (49). Thus, selection for closer linkage between the primary sex-determining factors and *SlBAM1* might have contributed to suppressing recombination in the most left-hand part of the ancestrally recombining region, forming Stratum 0.

It is unclear how or why the other strata evolved. If the ancestral state was like the present X, Stratum 1 formation involved cessation of recombination between Stratum 0 and the portion of the X recombining region that did not stop recombining when Stratum 0 formed, plus the pericentromeric region that is recombinationally inactive in females, which includes a region corresponding to parts of the *S. vulgaris* chromosome 3 (Fig. 4d). Undetected SA polymorphisms in this region could have favoured this recombination suppression, and/or the creation of Stratum 2. However, alternatives are possible. It was recently proposed that successive “lucky” inversions (or other changes) might become fixed among Y chromosomes by chance, extending the non-recombining region without being favored through involvement of sexually antagonistic polymorphisms. In one model, re-evolution of recombination is prevented by evolution of low expression of Y-linked alleles, accompanied by higher expression of their X counterparts, a form of dosage compensation (5). However, estimates of gene expression in young leaves of male and female *S. latifolia*, do not support the involvement of X chromosome dosage compensation. Using expression levels normalized so that autosomal values are similar in both sexes, the mean male values are above half those in females (fig. S19, table S8), confirming the previously reported partial compensation (16–18). On the other hand, compensation in the less degenerated Stratum 1 (which is probably still degenerating) appears similar to that in Stratum 0. In the younger Stratum 2, which is much less degenerated, expression is similar in both sexes (fig. S19), whereas the Lenormand and Roze (5) model predicts higher expression in females when Y-linked alleles are degenerating and being down-regulated, and their X alleles up-regulated. The expression estimates also do not reflect DNA methylation patterns (fig. S20), as has been proposed (50).

Figure 5 proposes a model for the evolution of the *S. latifolia* giant Y and X chromosomes. Establishment of the SPF mutation creating females, was followed by the appearance of the GSF (*SlCLV3*) in a closely linked genome region, creating a proto-X/Y pair (perhaps involving chromosome fusions). Selection for closer linkage led to suppressed recombination, creating an incipient Y. This was followed by Y-specific duplications, which would have helped to prevent recombination, to form Stratum 0. The non-recombining region thereafter accumulated LTR-TE insertions, which enlarged the Y-linked region. Fusions may have led to expansion of the rarely recombining X chromosome pericentromeric region, via crossover localization to regions near the ends of chromosomes; when a genome region is added to a chromosome end, part of the former recombinationally active region therefore probably becomes an inactive pericentromeric region, in which TE insertions start accumulating. The current Y chromosome is still undergoing rearrangements, probably partly due to TE activities, and stable heterochromatic epimarks, including DNA methylation are present throughout the MSY. Genetic degeneration, with many X-linked genes becoming hemizygous, is evident, and dosage compensation has started to evolve.

**Figure 5.**
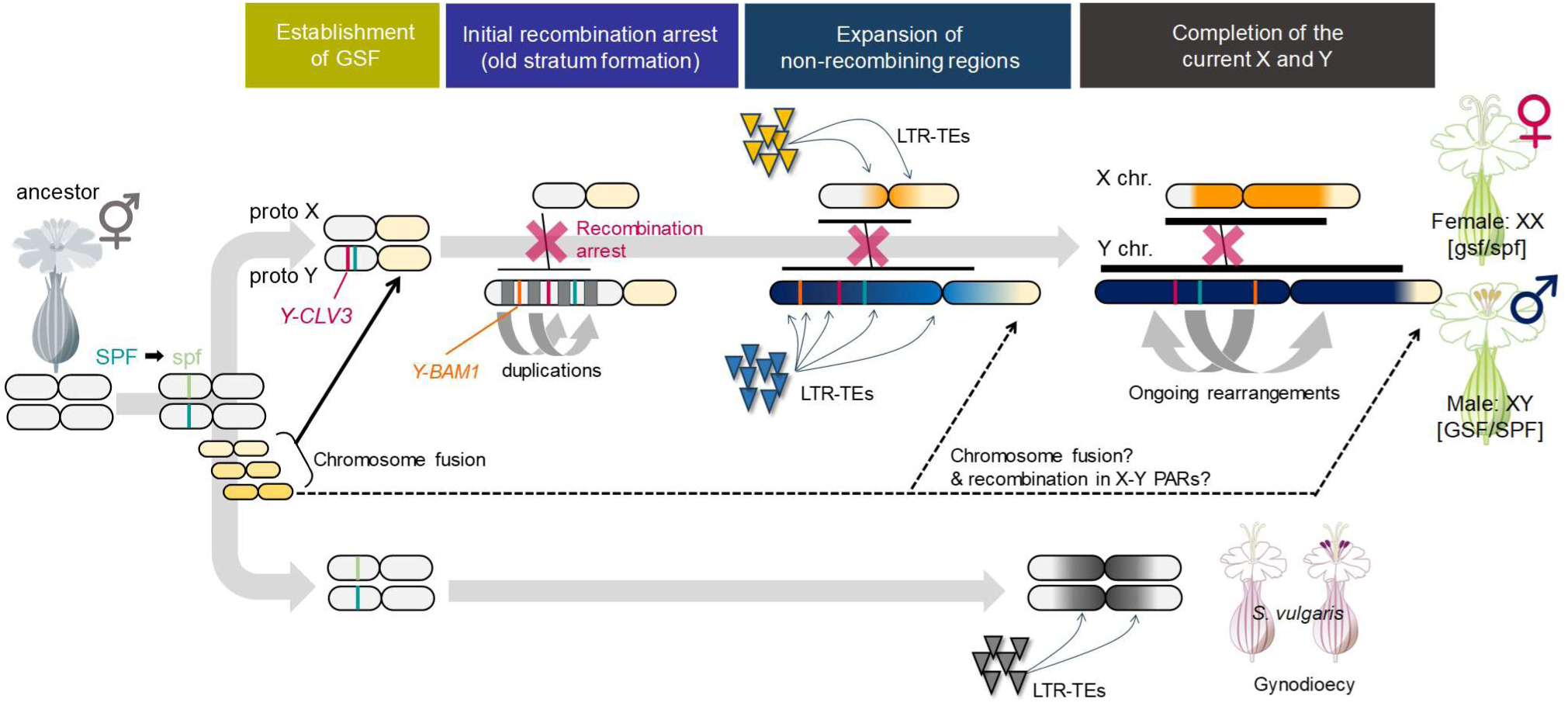
Model for the rapid and dynamic sex chromosome evolution in *S. latifolia*.

Although the *S. latifolia* XY pair is metacentric, with a single centromere, the lack of mature repeats in our CENH3-peak analyses suggests that the centromere may have shifted along with Y other chromosome rearrangements. Chromosome fusions may have been involved in formation of the present X/Y chromosome pair, and might also have occurred more recently, as suggested in Fig. 5. Our *S. latifolia* genome sequence helps to explain the evolution of a giant Y (and its structural rearrangements). We can also explain why the X chromosome is large. Its lack of structural rearrangements suggests that its large pericentromeric region does not completely lack recombination, but occasionally recombines. TE insertions, including recent female-specific proliferation of some active retrotransposon types (51), can therefore enlarge the region. However, even rare recombination will prevent gene loss, and degeneration therefore mainly affects the Y. Our results support the model involving selection for close linkage between two sex-determining factors that initially recombined. We also describe a Y-linked gene that controls a major sexual dimorphism in the species, strongly suggesting that that its alleles are sexually antagonistic. This polymorphism may also have contributed to selection for recombination suppression in the oldest stratum.

## Acknowledgements

We thank Dr. Isabelle M. Henry and Dr. Luca Comai (Dept. Plant Biol and Genome Center University of California Davis, USA) for discussion and comments for this study.

## Funding

This work was supported by PRESTO from Japan Science and Technology Agency (JST) [JPMJPR20D1] and Grant-in-Aid for Transformative Research Areas (A) from JSPS [22H05172 and 22H05173] to T.A., [22H05181] to K.S., and [23H04747] to Y.I.

## Authors contributions

Conceptualization: TA, DC

Methodology: TA, NF, KM, KS, YI, KU, DC

Investigation: TA, NF, KM, KS, KN, AH, EK, RK, MN, KU

Visualization: TA, NF, KM, KN, AH, KU

Funding acquisition: TA, NF, KS

Project administration: TA, DC

Supervision: TA, KU, DC

Writing – original draft: TA, DC

Writing – review & editing: TA, NF, KM, AH, KU, DC

## Competing interests

Authors declare that they have no competing interests.

## Data and materials availability

## Notes

### Competing Interest Statement

The authors have declared no competing interest.

