## Supplementary material for "Rapid and dynamic evolution of a giant Y chromosome in *Silene latifolia*": Methods and Supplementary Figures S1-S20

**Supplementary Figures S1-S20**

**Supplementary Tables S1-S10**

### Rapid and dynamic evolution of a giant Y chromosome in *Silene latifolia*


**Authors**

Takashi Akagi[1,2,*,§], Naoko Fujita[1,*], Kanae Masuda[1], Kenta Shirasawa[3], Kiyotaka Nagaki[4], Ayano Horiuchi[1], Eriko Kuwada[1], Riko Kunou[1], Koki Nakamura[4], Yoko Ikeda[4], Koichiro Ushijima[1], Deborah Charlesworth[5]

**Affiliations**

1. Graduate School of Environmental and Life Science, Okayama University, Okayama, Japan

2. Japan Science and Technology Agency (JST), PRESTO, Kawaguchi-shi, Saitama 332-0012, Japan

3. Kazusa DNA Research Institute, Kazusa-Kamatari, Kisarazu, Chiba, 292-0818, Japan

4. Institute of Plant Science and Resources, Okayama University, Kurashiki, Okayama, Japan

5. Institute of Ecology and Evolution, University of Edinburgh, Charlotte Auerbach Road, Edinburgh EH9 2FL, United Kingdom

*These authors contributed equally

§ Corresponding author

#### Methods

##### Plant materials

*Silene latifolia* seeds used in this study were the inbred lines from the previous study (1), which is defined as a German line in this study. *Silene vulgaris* seeds were purchased from *Jelitto* (Jelitto Staudensamen GmbH, Schwarmstedt, Germany). The seeds were surface sterilized and sowed on Murashige and Skoog (MS) media in a growth chamber under conditions of 23°C with a daylength of 16 hours. The plants were grown on MS media for a month (four to six leaves) and transplanted to soil (Kumiai Nippi Engei Baido No.1, Nihon Hiryo Co., Ltd., Tokyo, Japan) in a pot (Φ 9 cm x 7.5 cm). For *S. latifolia*, 228 individuals of an F1 population derived from the hybridization of two German lines, were used for the construction of a genetic map. For *S. vulgaris* plants, three individuals were selected as the parents according to their flower phenotypes (two hermaphrodites, namely SvPH1 and SvPH2, and one female, namely SvPF1). The hermaphroditic individual SvPH1, of which the genome was sequenced in this study, constantly produced hermaphroditic flowers. SvPH2 produced mostly hermaphroditic but occasionally female flowers. Their 45 self and 55 F1 hybrid progenies (SvPH2 x SvPH2 and SvPF1 x SvPH1, respectively) were used for ddRAD-sequencing for the construction of a genetic map. The details of plant samples are summarized in table S9.

##### Genome assembly

Genomic DNAs were extracted from young leaves of male *S. latifolia* (a German line) and hermaphrodite *S. vulgaris* (SvPH1) Genomic DNA was extracted from young leaves using NucleoSpin Plant II kit (Takara Bio Inc., Shiga, Japan) or QIAGEN Genomic-tip 20/G (Qiagen, Hilden, Germany). According to the previous report (2), the genome DNA was sheared into ~20 kb fragments with a g-tube machine (Covaris, Woburn, MA, USA) to construct a HiFi SMRTbell library with a SMRTbell Express Template Prep Kit 2.0 (PacBio, Menlo Park, CA, USA). The library DNA was fractionated using the BluePippin (Sage Science, Beverly, MA, USA) to eliminate small fragments and sequenced with a Four SMRT cell 8 M on the Sequel II system (PacBio). The sequence reads were converted into HiFi reads with the CCS pipelines (PacBio, https://ccs.how) and assembled with Hifiasm (3) to generate initial assembled contigs.

For further contig anchoring, we applied Bionano optical mapping and anchoring with genetic maps. The genomic DNAs were labeled with the DLE-1 enzyme and subjected to the optical mapping bb Bionano Saphyr system (Bionano Genomics, San Diego, CA, USA). The initial assembled contigs were scaffolded by hybrid assembly method of Bionano Solve package (available at https://bionanogenomics.com/support/software-downloads/), and generated SuperScaffold contigs. We constructed the genetic maps from ddRAD-seq date of the described F1 populations of *S. latifolia* and *S. vulgaris* to anchor SuperScaffold contigs and generate pseudomolecules. The ddRAD-seq reads were aligned to the SuperScaffolds by minimap2 (4) and single nucleotide polymorphisms (SNPs) were called by using varscan2 (5). After filtration of minor SNPs, linkage map was constructed

by Lep-Map3 (6). The linkage map was then used to determine the order and orientations of the SuperScaffolds on each chromosome by Allmaps (7), except for Y chromosome of *S. latifolia*. The linkage map was further used to determine the genetic recombination rate by MareyMap online (https://lbbe-shiny.univ-lyon1.fr/MareyMapOnline/).
Intragenomic collinearity (or paleoduplication event) was evaluated by all-versus-all BLASTP analyses (<$e^{-50}$ in blastp), for the whole genes, followed by the selection with threshold values of silent divergence (*dS*; described later) and visualization with the Circa software (https://omgenomics.com/circa/).

**Cytogenetic analyses**

Chromosomes were prepared from flower buds of *S. latifolia*. The flower buds were treated in 50 μM colchicine at 4ºC for 4 h, and then kept in ice-cold water for 24 h. The treated buds were fixed in ethanol:acetic acid=3:1. The fixed buds were washed in distilled water twice and digested in an enzyme mix containing 1% (w/v) cellulase Onozuka RS (Yakult Pharmaceutical Industry Co., Ltd., Tokyo, Japan), 0.5% (w/v) pectoriase Y-23 (Kikkoman Biochemifa Company, Tokyo, Japan) and 1% (w/v) cytohelicase (C8274; Sigma-Aldrich, St. Louis, MO, US) for 1 h. The digested buds were washed in distilled water twice and re-fixed in methanol:acetic acid=3:1 and suspended in a small amount of the fixative. The suspended cells were spreaded by a flame-dry method. Chromosomes of the spread cells were counterstained with 4,6-diamino-2-phenylindole (DAPI). The stained chromosomes were captured using a chilled charge-coupled device (CCD) camera, AxioCam HR (Carl Zeiss, Oberkochen, Germany), and images were pseudo-colored and processed using Zen software (Carl Zeiss).
A set of myTags probes specific for the Y-chromosome was designed and synthesized by Arbor Biosciences. FISH analysis was performed as previously described (8). The synthesized probes were hybridized to the spread chromosomes. After washing in 2 x SSC at RT, chromosomes were counterstained DAPI. The stained chromosomes and myTags signals were captured and visualized using the system described above.
Based on the deduced *S. latifolia* CENH3 (g19732.t1) amino acid sequence, a peptide corresponding to the N-terminus of *S. latifolia* CENH3 (H2N- RIKQVATKSSASKRRSYASA -COOH) was synthesized and injected into two rabbits. The raised antisera were purified using an affinity Sepharose column consisting of the aforementioned peptide. Immunostaining was conducted as previously described (9) with modifications. In brief, flower buds were fixed in phosphate-buffered saline (PBS) containing 3% (w/v) paraformaldehyde and 0.2% (v/v) Triton X-100. The fixed buds were washed and digested with a mixture of 1% (w/v) cellulase Onozuka RS (Yakult Pharmaceutical Industry, Tokyo, Japan) and 0.5% (w/v) pectolyase Y-23 (Seishin Pharmaceuticals, Tokyo, Japan) and then squashed onto slides coated with poly-L-lysine (Matsunami, Osaka, Japan). A 1:100 dilution of the purified anti-CENH3 antibody was applied to the slides. The antibody was detected using 1:400 dilutions of Alexa Fluor 555-labeled anti-rabbit antibodies (Molecular Probes, OR, USA). Chromosomes were counterstained with DAPI.

Immunosignals and stained chromosomes were captured and visualized using the system described above.

**Repeat and gene annotations**

Repetitive sequences in the assemblies were identified with phRAIDER (10), according to the previous report (2). In this study, repetitive sequences were categorized into nine types: short interspersed nuclear elements (SINEs), long interspersed nuclear elements (LINEs), long terminal repeat (LTR) elements, DNA elements, small RNA, satellites, simple repeats, low complexity repeats, and unclassified.

Protein-coding genes were annotated in the genomic sequence of which repetitive sequences detected above were masked. Gene prediction was conducted with the Braker2 pipeline, trained with Illumina short reads mRNA-seq data from a variety of organs and developmental processes (see mRNA-seq datasets deposited to DDBJ; PRJDB16402 for the bioproject), according to the previous pipeline (11). The completeness of the assemblies was assessed using the BUSCO score (12).

**DNA methylome library**

DNA methylome libraries were constructed with genomic DNA extracted from young leaves ($5^{th}$ leaves from 4 biological replicates for each male and female sibling), according to previous reports (13, 14). gDNA was fragmented to approx. 300-400bp segments using Picoruptor sonicator (Diagenode), followed by end-repairing and adenosine-tailing steps to ligate with a cytosine-methylated adapter, by using KAPA HyperPlus Kit (KAPA Biosystems). The adapter-ligated DNA fragments were purified with AMPure XP beads (Beckman Coulter), and then subjected to a bisulfite conversion step with the EZ DNA Methylation-Gold kit (Zymo Research, USA). The bisulfite-converted DNA fragments were enriched by 6-8 cycles of PCR, and then purified using AMPure.

**ChIP-Seq library**

Total 0.15g leaf of a male *S. latifolia*, used for the whole genome assembly, was frozen with liquid nitrogen, ground into fine powder, and crosslinked with 1% formaldehyde for 10 min at room temperature. Crosslinking was quenched by the addition of glycine to a final concentration of 125 mM. Nuclear-extraction and immunoprecipitation was carried out with modifications of Nozawa et al. 2022 (15). The chromatin precipitate was resuspended in 150 µL SDS lysis buffer (50 mM Tris-HCl pH 8.0, 10 mM EDTA, 1% SDS) and sheared using Picoruptor sonicator (Diagenode) for 12 cycles of 30 s “ON” and 30 s “OFF” at 10 °C. After centrifugation at 13,000 × g at 4°C for 10 min, 100 µL of the supernatant was used for immunoprecipitation. Immunoprecipitation was performed with 3 µg of *S. latifolia* anti-CENH3 antibody (described above), anti-H3K9me2 antibody (mAbcam1220; Abcam), or anti-H3 antibody (ab1791; Abcam). After washing, immune complexes were eluted with elution buffer (10 mM Tris–HCl, pH 8.0, 0.3 M NaCl, 5 mM EDTA, 0.5% SDS) and DNA was reverse-crosslinked by incubating overnight at 65 °C. After proteinase K and RNase treatment, DNA was purified by phenol-chloroform method. Total 1.5–2ng of DNA was used for ChIP-seq library preparation with the KAPA HyperPlus Kit (KAPA Biosystems).

**Illumina reads sequencing and processing for DNA methylome and ChIP-seq libraries**

The Illumina libraries were sequenced using Illumina's HiSeqX (150-bp paired-end reads). All Illumina sequencing was conducted at Macrogen Japan. Raw Illumina reads were processed using custom Python scripts developed in the Comai laboratory and available online (http://comailab.genomecenter.ucdavis.edu/index.php/), as previously described (16).

For DNA methylome data, mapping and methylated-cytosine detection were conducted with the methylpy pipeline (https://github.com/yupenghe/methylpy, 17). The weighted methylation level (18) was calculated separately for CG, CHG, and CHH contexts. The averaged methylation values surrounding the gene bodies were visualized with deeptools2 (19).

For ChIP-Seq data, the preprocessed reads were aligned to the whole chromosomes of *S. latifolia*, with Burrows-Wheeler Aligner (BWA)-mem (20), with the default parameters. The reads numbers mapped to each 1-Mb bin were integrated by using a custom Python script, bin-by-sam.py (21, 22), to estimate CENH3 and H3K9me2 signals, with standardization by H3 amount.

**Centromeric repeat assessment**

Approximately 5Mb genomic regions surrounding putative CENH3-peak regions were extracted from each chromosome. Except chromosome 10, we could detect clear signals of *S. latifolia* CENH3. StainedGlass package (23) was used to generate sequence identity heat maps, and the analysis was conducted using a 2-kb window with the mm_f option set to 10 kb.

***de novo* transposable elements (TEs) annotations and phylogenetic analysis**

TEs in the genomes of the two *Silene* species (*S. latifolia* and *S. vulgaris*) were analyzed with the Extensive de novo TE Annotator pipeline (24), which integrates structure-based and homology-based approaches for TE identification, including LTRharvest (v.1.5.10) (25), LTR_FINDER_parallel (v.1.0) (26, 27), LTR_retriever (v.2.6) (28), TIR-Learner (v.1.23) (29), Generic Repeat Finder (v.1.0) (30), HelitronScanner (v.1.0) (31) and TEsorter (32), with extra basic and advanced filters. Clustering of TEs within the Gypsy and Unknown family on chr1-4 in *S. vulgaris* and on chrX-Y in *S. latifolia* was conducted using cd-hit-est (33), with the c = 0.8 (>80% sequence identity) option. The TEs counts clustered with higher identity (>80% sequence identity, TE counts > 100) were visualized using ComplexHeatmap (34). These TEs were subclustered using cd-hit-est, with the c = 0.85, 0.875, 0.9, 0.925, 0.95, and 0.975 to calculate the diversification of TEs. For the evolutionary topology of TEs within clusters, their sequences (sequence length < 2,000 bp) were aligned with MAFFT v.7 (35) with the L-INS-i model (--maxiterate 1000 --adjustdirection), followed by automatic pruning using TrimAl (36). The alignments were used to construct phylogenic trees with the maximum likelihood method using IQ-TREE with 1,000 bootstraps (37). Phylogenetic trees were visualized using ggtree (38). The estimated silent divergence (*dS*) values (used as the scale bars in Fig. 4) were calculated based on in-codon frame alignments of the protein-coding sequences, in some full-length TEs, as described below.

**Synteny analyses between X and Y, and between *S. latifolia* and *S. vulgaris***

Sequence-based synteny analysis was performed using minimap2 (4) and visualized with dotPlotly (https://github.com/tpoorten/dotPlotly), with a minimum alignment length of 50,000 bp. Large scale synteny based on the gene orders was examined with MCScanX (39), in which the detected collinearity was visualized using SynVisio (https://synvisio.github.io/#/). All-versus-all BLASTP analyses were performed among the protein sequences in the *S. latifolia* X and Y, or in the *S. latifolia* and *S. vulgaris* whole genomes, with an e-value cut-off of <1e$^{-10}$ for the comparison of X and Y, and <1e$^{-20}$ for the intragenomic comparison of *S. latifolia* and *S. vulgaris*. Syntenic blocks were defined by using MCScanX, with BLASTP results and gff data. Collinearity of gene orders with specific *dS* values was analyzed and visualized with SynMap and GEvo in CoGe (40, 41, https://genomevolution.org/coge/).

**Detection of genetic diversity and positive selections**

Genetic diversity among the orthologous or paralogous gene pairs, which exhibited significant sequence similarity (<e$^{-10}$ in blastp), was detected based on an in-codon frame alignment using Pal2Nal and MAFFT ver. 7 under the L-INS-i model (35), according to the previous method (2). The alignments were analyzed with MEGA X (42) to estimate the Jukes and Cantor corrected values of silent divergence (*dS*). For transition of *dS* values according to the genetic map of X chromosome, change-point detection was performed with Pettitt's test, by using "pettitt.test" function in the "trend" package of R.

Evolutionary rate, or *dN/dS* value, was calculated from the described in-codon frame alignments, using non-synonymous substitution (*dN*) and synonymous substitution (*dS*) rates detected by MEGA X.

McDonald and Kreitman test for inspection of neutral selection, was conducted with in-codon frame alignments of *S. latifolia* Y chromosomal paralogues, which was putatively derived from Y-specific duplication events, and their orthologues in X chromosome and in *S. vulgaris*, by using DnaSP v.6 (43).

Neutral selection on the Y chromosomal paralogues (defined as "species 1" in DnaSP) was tested by using *S. latifolia* X-alleles and *S. vulgaris* orthologues as the outgroup (defined as "species 2"). Neutral index (NI) and p-values with G-test were calculated as the indexes of neutral selection.

Y-lost index, which was defined as the rate of X-specific genes (or Y-lost genes) in the category of the genes conserved between *S. latifolia* and *S. vulgaris*, was evaluated with two criteria: (i) complete loss and (ii) annotation loss. Annotation loss is the situation that Y chromosome has no annotated protein-coding gene homologous to an X gene, in blastp analyses (<$e^{-10}$ for the threshold). Complete loss is the situation that Y genomic sequences include no homologous sequences to an X gene. For detection of complete loss, tblastn analysis (<$e^{-10}$ for the threshold) was conducted with X genes as the queries and whole genome sequences as the database. If an autosomal fragment exhibits the highest hit (or no one hits) against an X gene, we counted this as a Y-lost in the complete loss criterion.

**Detection of dosage compensation**

For the transcriptome analyses, mRNA-seq libraries were constructed with RNAs extracted from young leaves (5$^{th}$ leaves from 4 biological replicates for each male and female sibling), according to previous reports (44). To examine the effects of DNA methylation status for dosage compensation, we collected half of the leaf for RNA and the other half for DNA methylome analyses. All sampling was performed at midday. Total RNA was isolated using PureLink Plant RNA Reagent (Invitrogen). mRNA was purified using the Dynabeads mRNA purification kit (Life Technologies). mRNA-seq library was synthesized via KAPA RNA HyperPrep kit (Kapa Bioscience), followed by a DNA cleanup step with AMPure XP beads (Beckman Coulter; AMPure:reaction, 0.7:1), according to the previous report (44). The resultant libraries were sequenced on Illumina's HiSeq X sequencer (150-bp single-end reads). All Illumina sequencing was conducted at Macrogen Japan. Sequencing reads were processed using Python scripts (https://github.com/Comai-Lab/allprep/blob/master/allprep-13.py), as described. The mRNA-seq reads were mapped to the whole coding regions in *S. latifolia*, using the Burrows-Wheeler aligner (BWA)-mem (http://bio-bwa.sourceforge.net/) with the default parameters. We preliminarily checked that bwa-mem with the default mapping parameters could distinguish most of the X and Y alleles. Mapped read numbers were counted from the aligned SAM files using custom R scripts, according to the previous report (16). For the selection of X-specific (or X hemyzygous) genes, the “annotation loss” category in the Y-lost index described above, was applied. Standardized expression levels (read numbers per kilobase and millions reads: RPKM) were calculated in each male and female, and their biases were plotted in the X-genetic map.

**Virus induced gene silencing with ALSV in *S. latifolia***

Virus induced gene silencing (VIGS) with apple latent spherical virus (ALSV) vector was performed as previously described (1). Briefly, Y-SlCLV3-ALSV vector containing 204 bp fragment of *Y-SlCLV3* (shown in Figure S9) was prepared by introducing the synthesized DNA fragment (Azenda Life Sciences, MA, US) to SalI-BamHI site of the ALSV vector. The resulting Y-SlCLV3-ALSV vector, which was initially transformed into *Agrobacterium tumefaciens* EHA105, was inoculated into *Nicotiana benthamiana* plants by the agro-infiltration method. Viral RNAs were purified from the infected *N. benthamiana* plants and was precipitated onto gold particles. The viral RNA coated on gold particles were bombarded into the cotyledons of *S. latifolia* seedlings in an emergence stage by the particle bombardment GDS-80 (Nepa Gene Co., Ltd., Chiba, Japan).

**Transformation of *Nicotiana tabacum***

Genomic sequences of the Y-encoded *SlBAM1* (*Y-BAM1*), including approx. 2-kb 5' promoter region, were amplified by PCR using PrimeSTAR Max (TaKaRa) from a male *S. latifolia* (a German line grown in Okayama University) gDNA extracted from young leaves (table S10 for primer information). The PCR amplicon was cloned into the pPLV02 vector (45) to place the genes under the native promoter, to design *pYBAM1::YBAM1* construct, using the InFusion (TaKaRa, Japan) pipeline, according to the previous report (16). Tobacco plants (*N. tabacum*) cv. Petit Havana SR1 were grown *in vitro* under white light with 16-h-light and 8-h-dark cycles at 24°C. The *pYBAM1::YBAM1* was introduced into the *A. tumefaciens* strain EHA105. Young leaves of tobacco plants were transformed according to the method as previously described (46, 47). Transgenic plants were selected on Murashige and Skoog (MS) medium supplemented with 100 µg/mL kanamycin.

**Sequence data deposition**

All of the genome sequences and the annotated data were deposited to Plant GARDEN (https://plantgarden.jp/en/index). The raw sequencing data have been deposited in the DDBJ database: Sequence Read Archives database (BioProject ID PRJDB16402, Run ID DRR496111-DRR496477 for genome sequencing, DNA methylome and ChIP-seq data, and BioProject ID PRJDB16382, Run ID SAMD00635100-00635106 and SAMD00636369-00636376 for RNA-seq data in flower buds.

**Figure S1**

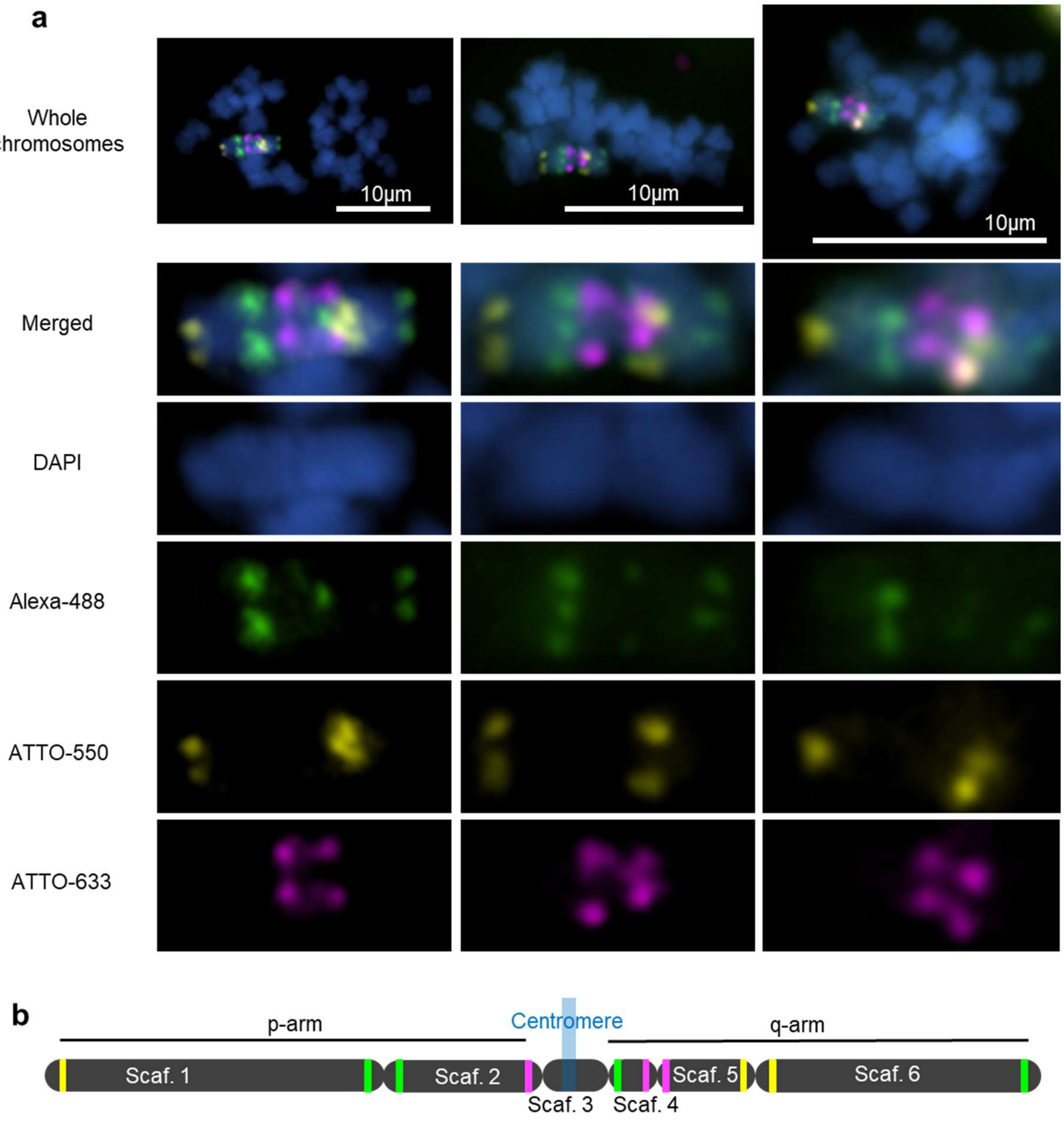


**Figure S1 Cytogenetic anchoring of the Y chromosomal contigs with FISH probes**

**a**, FISH anchoring of the genomic contigs with the myTags probes. **b**, The estimated structure of the Y chromosome, based on the FISH probe signals. The myTag probes were designed in MSY, except the right-end of the Scaffold 6, which is in PAR while targets the repetitive sequences specific to Y chromosome.

**Figure S2**

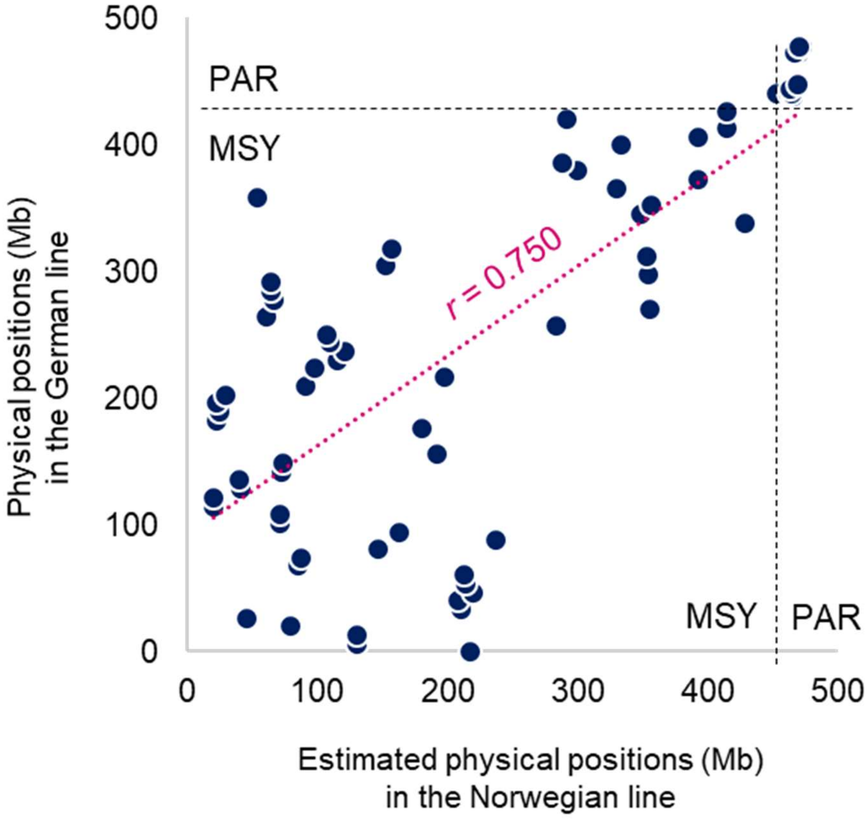


**Figure S2 Correlation of the genetic marker order in this study (x axis) with the positions suggested by deletion mapping (y axis, from Kazama et al. 2016 (48))**

Spearman's correlation test between the approximately estimated position of the genetic markers used in Kazama et al. 2016 (48) in the Norwegian line (K-line) (X-axis) and the physical position of them in our Y chromosome assembly in the German line (Y-axis). Although the correlation value was statistically high ($r$ = 0.75), the marker orders were highly rearranged between these two accessions.

**Fig S3.**

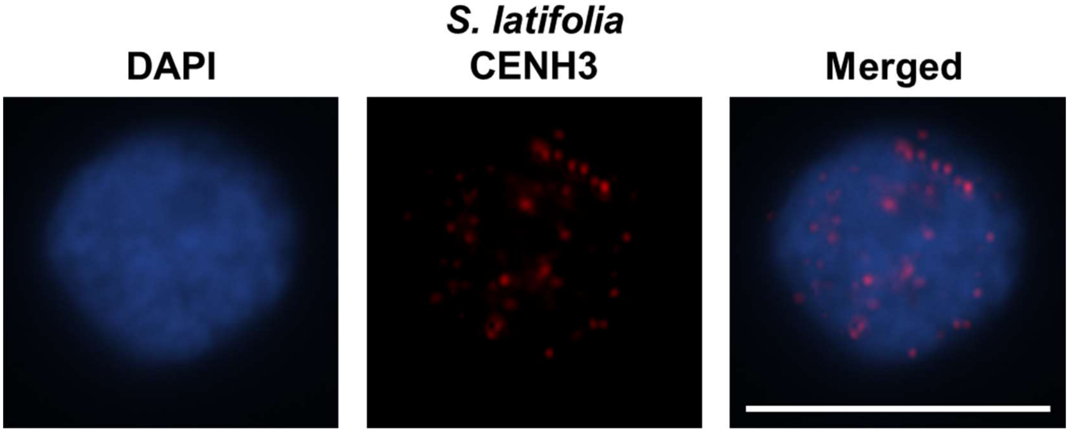


**Figure S3 Immunostaining with the anti-CENH3 antibody against a *S. latifolia* pro-metaphase cell.**

Cytogenetic observation of the custom anti-CENH3 antibody localization in the putative centromeric regions in pro-metaphase cells. Scale bar: 10 μm.

#### Figure S4

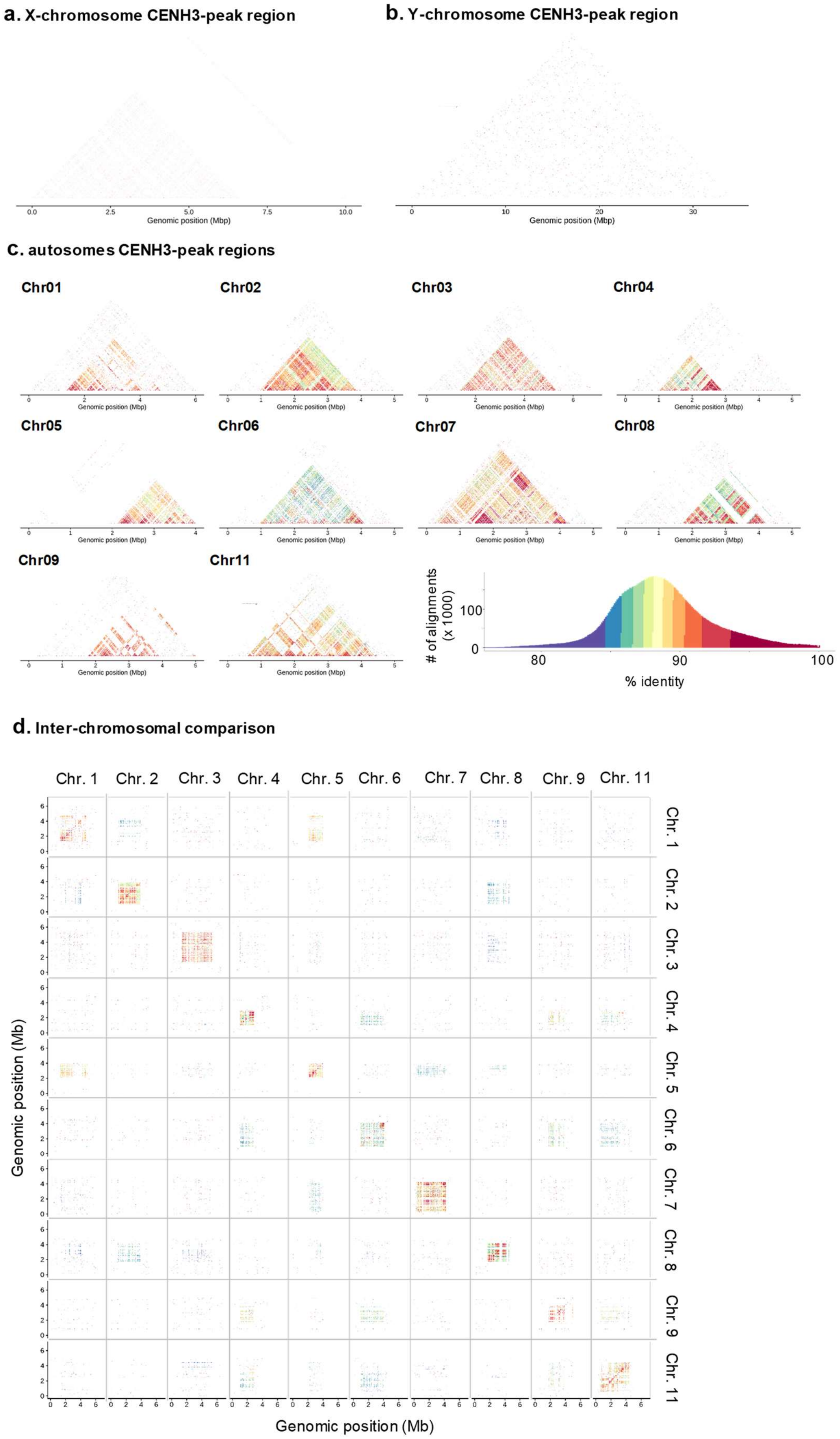

a. X-chromosome CENH3-peak region
0.0
2.5
5.0
7.5
10.0
Genomic position (Mbp)
b. Y-chromosome CENH3-peak region
0
10
20
30
Genomic position (Mbp)
c. autosomes CENH3-peak regions
Chr01
Chr02
Chr03
Chr04
Chr05
Chr06
Chr07
Chr08
Chr09
Chr11
Genomic position (Mbp)
### of alignments (x 1000)
100
0
80
90
100
% identity
d. Inter-chromosomal comparison
Chr. 1
Chr. 2
Chr. 3
Chr. 4
Chr. 5
Chr. 6
Chr. 7
Chr. 8
Chr. 9
Chr. 11
Genomic position (Mb)
Genomic position (Mb)

**Figure S4 Assessment of centromeric repeats in *S. latifolia***

Tandem repeats in self-synteny analyses surrounding the CENH3 ChIP-Seq peaks were visualized with heatmaps by using StainedGlass (23). X-chromosome CENH3-peak region formed faint tandem repeats (**a**), whereas no matured tandem repeats were detected in the Y chromosome CENH3-peak region (**b**). In comparison to the autosomes (**c**), the centromeric tandem repeats in the sex chromosomes were considerably immature. These implies frequent turnovers of the centromeric regions, especially in Y chromosome, which is consistent with drastic rearrangements of the marker orders even in a species (Fig. 2). **d**, Inter-chromosomal comparison of the tandem repeats in the centromeric regions. Not perfectly consistent with the case of Arabidopsis (49), centromeric repeats are not highly conserved amongst the chromosomes, and are evolved in a chromosome-specific manner.

#### Figure S5

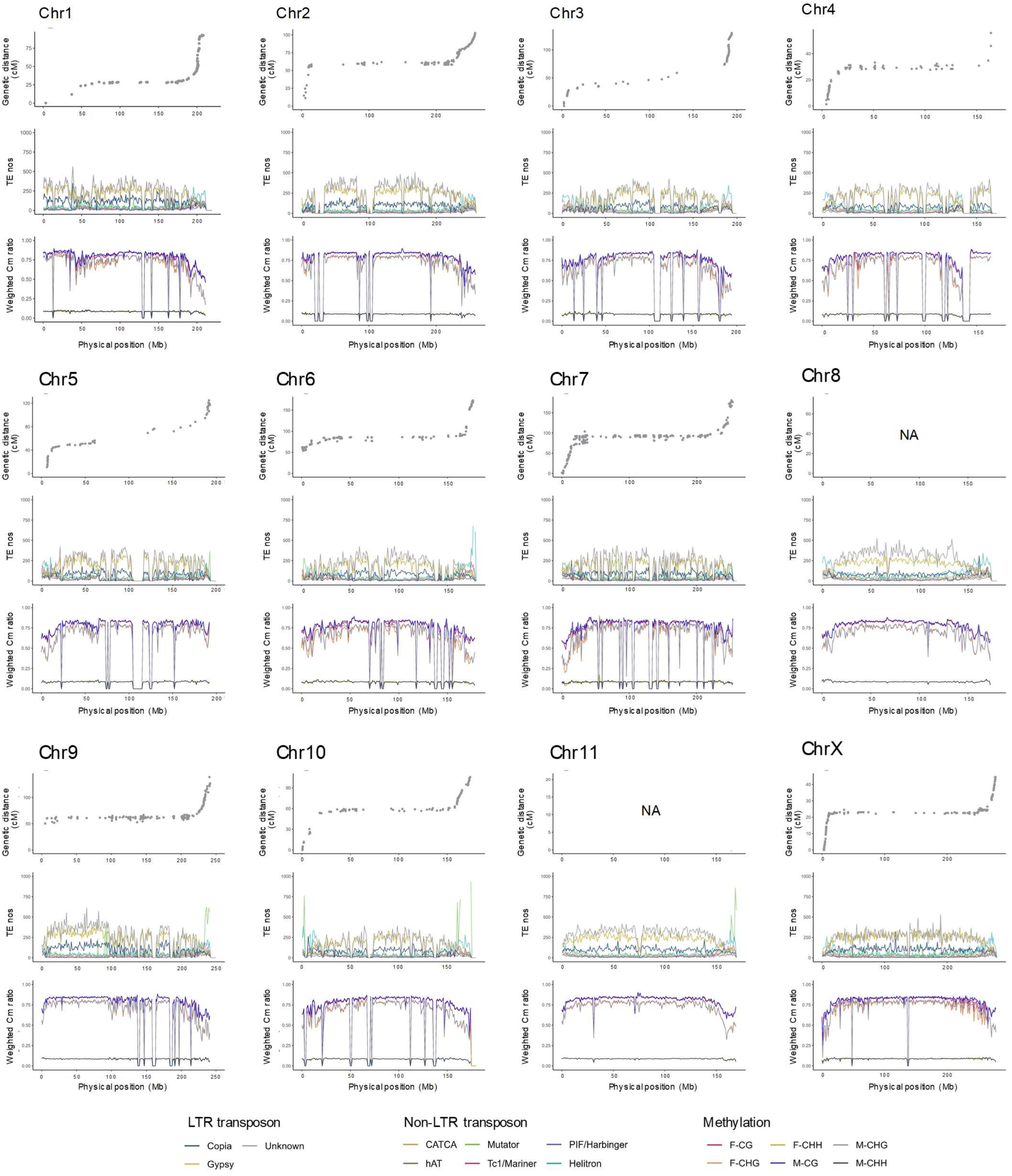


**Figure S5 Genetic recombination rates, distribution of transposons, and DNA methylation in *S. latifolia*.**

Genetic distances according to the physical positions (upper row), TE distributions (middle row), and

DNA methylation levels (in leaves of males and females, separately) (bottom row) were given for 11 autosomes and X-chromosome. For TEs, nine classes were separately provided per 1Mb bin. For DNA methylation levels, the three contexts (CG, CHG, and CHH) were separately provided for both females and males (F-CG, F-CHG, F-CHH for females, and M-CG, M-CHG, M-CHH for males, respectively). The long recombination arrested regions, mostly in the middle area of each chromosome, were highly enriched with LTR-type TEs, especially *Gypsy* and *unknown* classes, and with CG/CHG DNA methylation.

**Figure S6**

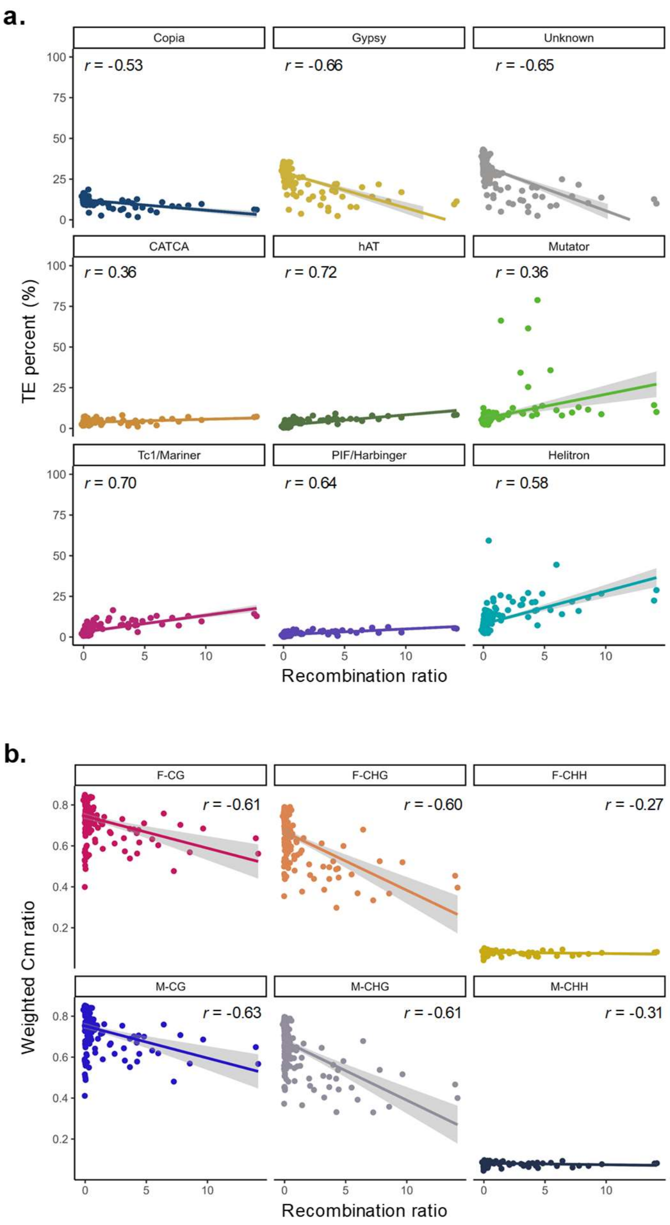


**Figure S6 Correlations between densities of different TE classes/DNA methylation levels and recombination rates in *S. latifolia***

The Pearson's moment correlation test was performed between the recombination rate (per 10Mb) and each TE class accumulation (**a**) or DNA methylation level (**b**). Consistent with the trends observed in fig. S5, the recombination rates were negatively correlated to LTR-type TEs (or *Copia*, *Gypsy*, and *unknown* classes) and CG/CHG DNA methylations.

#### Figure S7

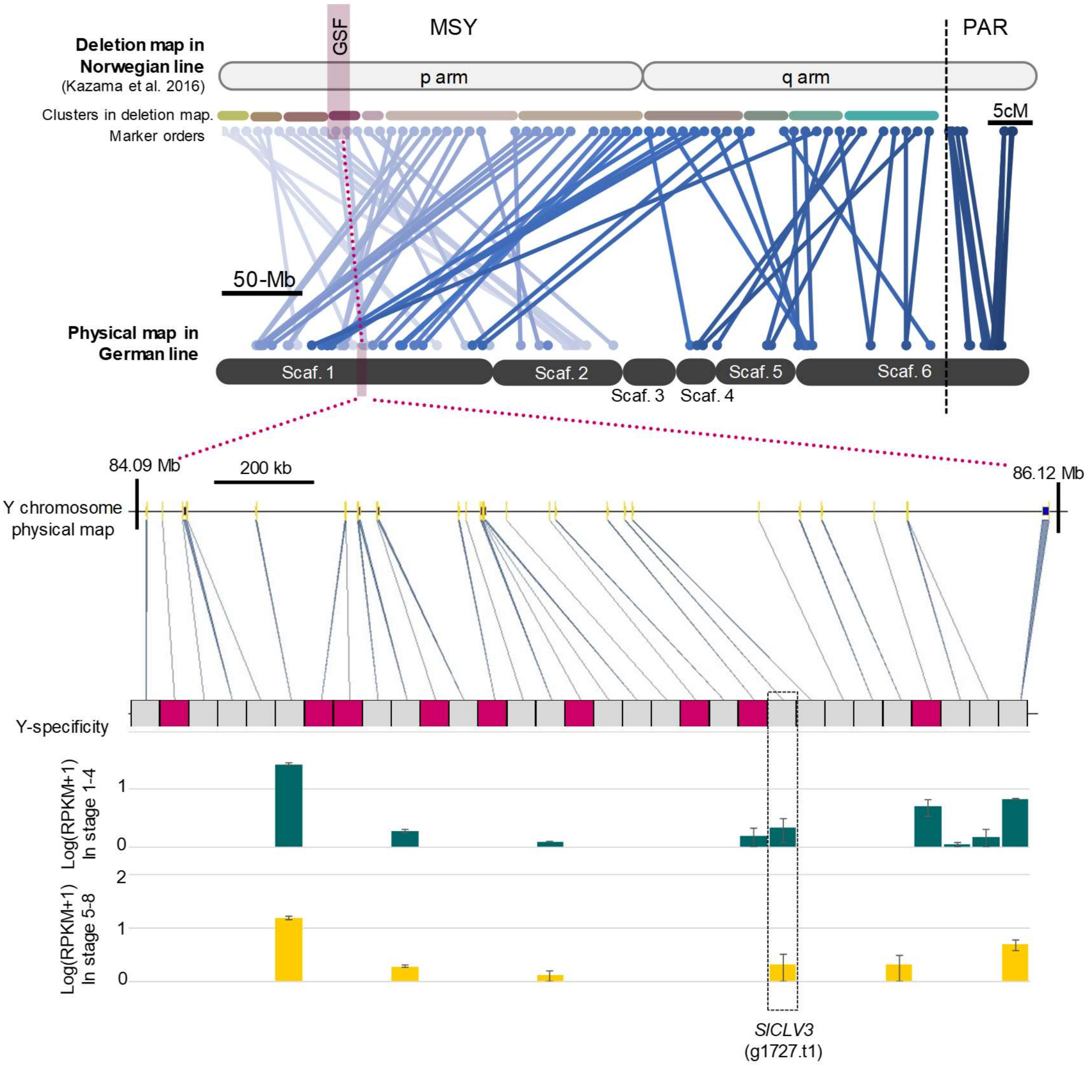


**Figure S7 Characterization of the Y-p arm region that is inferred by deletion mapping to include the GSF, and expression of potential candidate genes located in the nearby region**

Based on the deletion mapping by Kazama et al. 2016 (48), we could define GSF-included approx. 2.03Mb region. This region includes 31 canonical genes, of which nine were Y-specific. Integration of transcriptomic data (in early and late flower primordia differentiation stages, defined as stage 1-4 and 5-7, respectively,1) indicated only 10 genes were substantially expressed (RPKM > 1.0 in either of the stages).

**Figure S8**

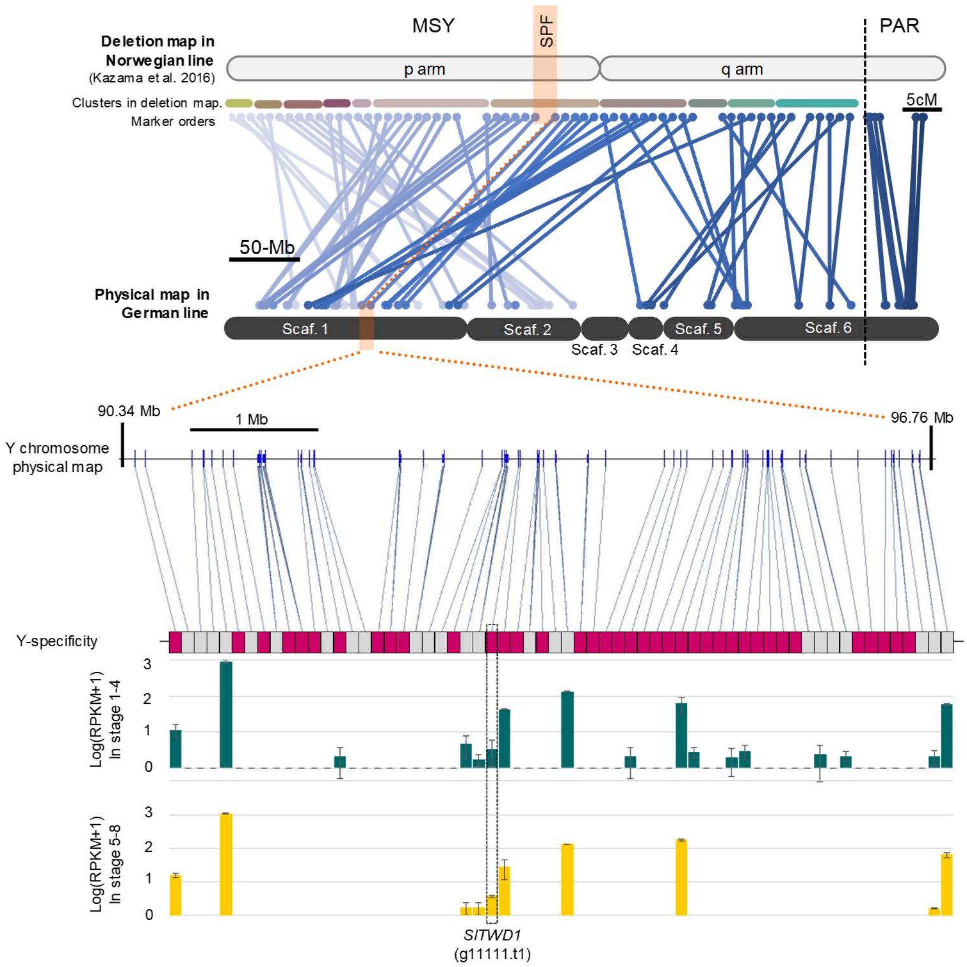


**Figure S8 Characterization of the Y-p arm region that is inferred by deletion mapping to include the SPF, and expression of potential candidate genes in the region**

Based on the deletion mapping by Kazama et al. 2016 (48), we could define SPF-included approx. 6.42Mb region. This region includes 62 canonical genes, of which 37 were Y-specific. Integration of transcriptomic data (in early and late flower primordia differentiation stages, defined as stage 1-4 and 5-7, respectively, 1) indicated 17 genes were substantially expressed (RPKM > 1.0 in either of the stages).

**Figure S9**

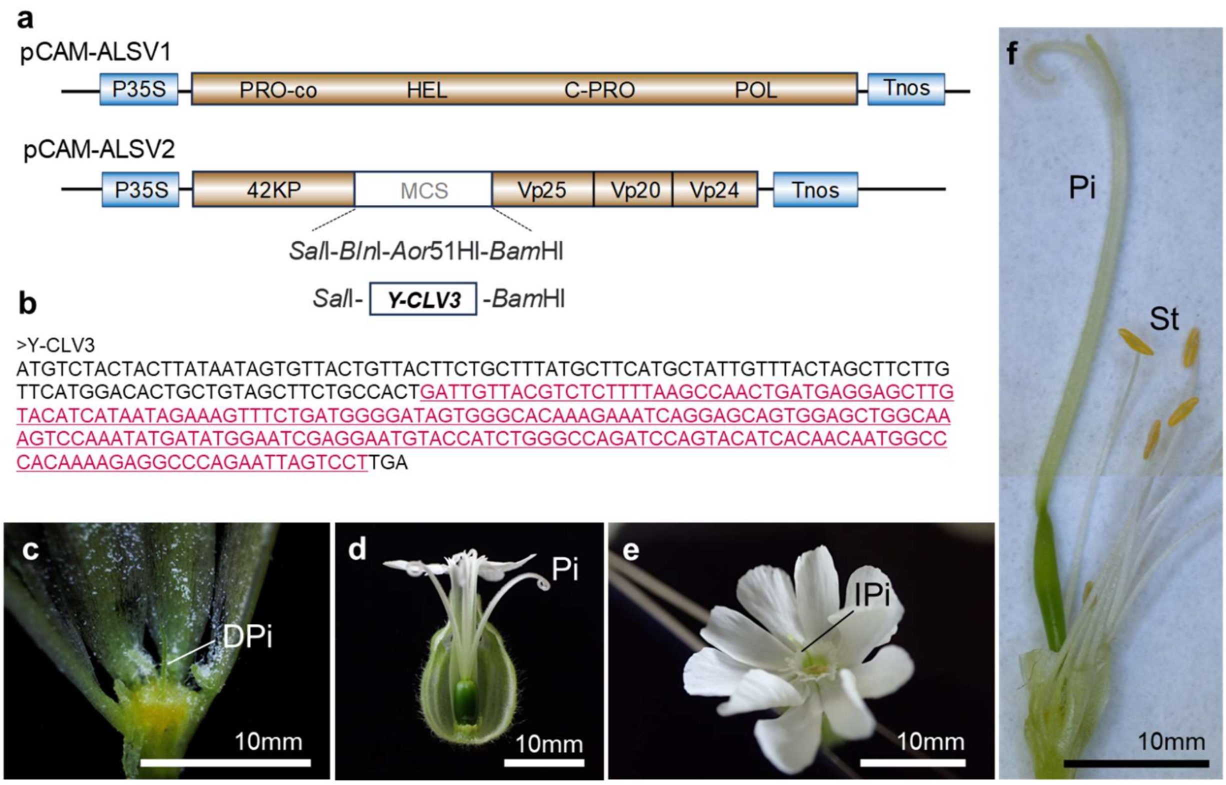


**Figure S9 Evidence supporting ALSV-induced VIGS targeting of the *Y-SlCLV3* gene**

**a**, Constructs for the ALSV vector inserted with *Y-SlCLV3* sequences (given in **b** for the inserted sequences highlighted in magenta and underline). Control male (**c**) has defected pistil (DPi), while control female (**d**) shows long pistil (Pi). The males infected with ALSV targeting *Y-SlCLV3*, exhibited imperfect pistil (IPi) (**e**) or fertile long pistil (**f**), with fertile stamens (St).

**Figure S10**

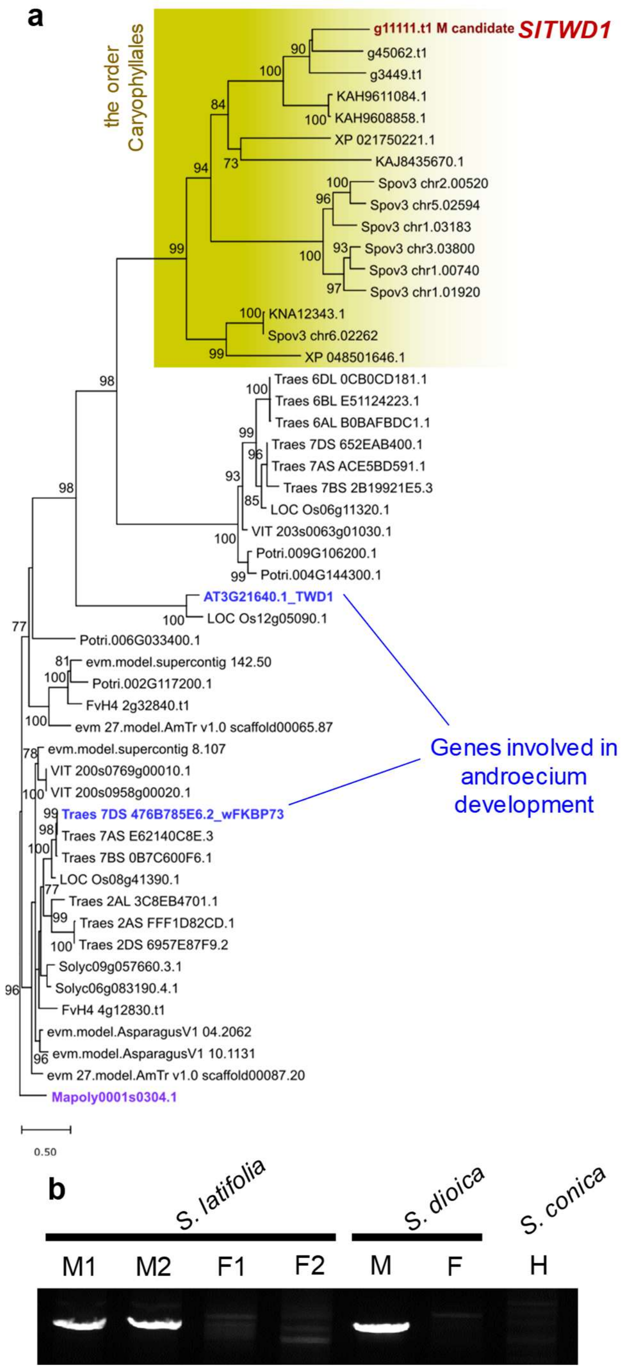


**Figure S10 Characterization of the *SlTWD1* gene as a candidate for the *S. latifolia* SPF (or M factor)**

**a**, Maximum likelihood (ML) phylogenetic trees of SlTWD1. For the OTU prefixes, AT: Arabidopsis, LOC: *Oryza sativa*, Traes: *Triticum aestivum*, Solyc: *Solanum lycopersicum*, VIT: *Vitis vinifera*, FvH4: *Fragaria vesca*, Potri: *Populus trichocarpa*, evm.model.Asparagus: *Asparagus officinalis*, Spov3: *Spinacia oleracea*. The genes highlighted in blue can act for stamen development (50, 51). The closest orthologue in liverwort (*Marchantia polymorpha*) was set as the outgroup, highlighted in purple.

**b**, PCR amplification of *SlTWD1*, in dioecious *S. latifolia* and *S. dioica*, and hermaphrodite *S. conica*, exhibited male-specific signals (M: male, F: female, and H: hermaphrodite). Note that *SlTWD1*-like sequences are highly repetitive in the genome, and often non-specific signals are amplified.

**Figure S11**

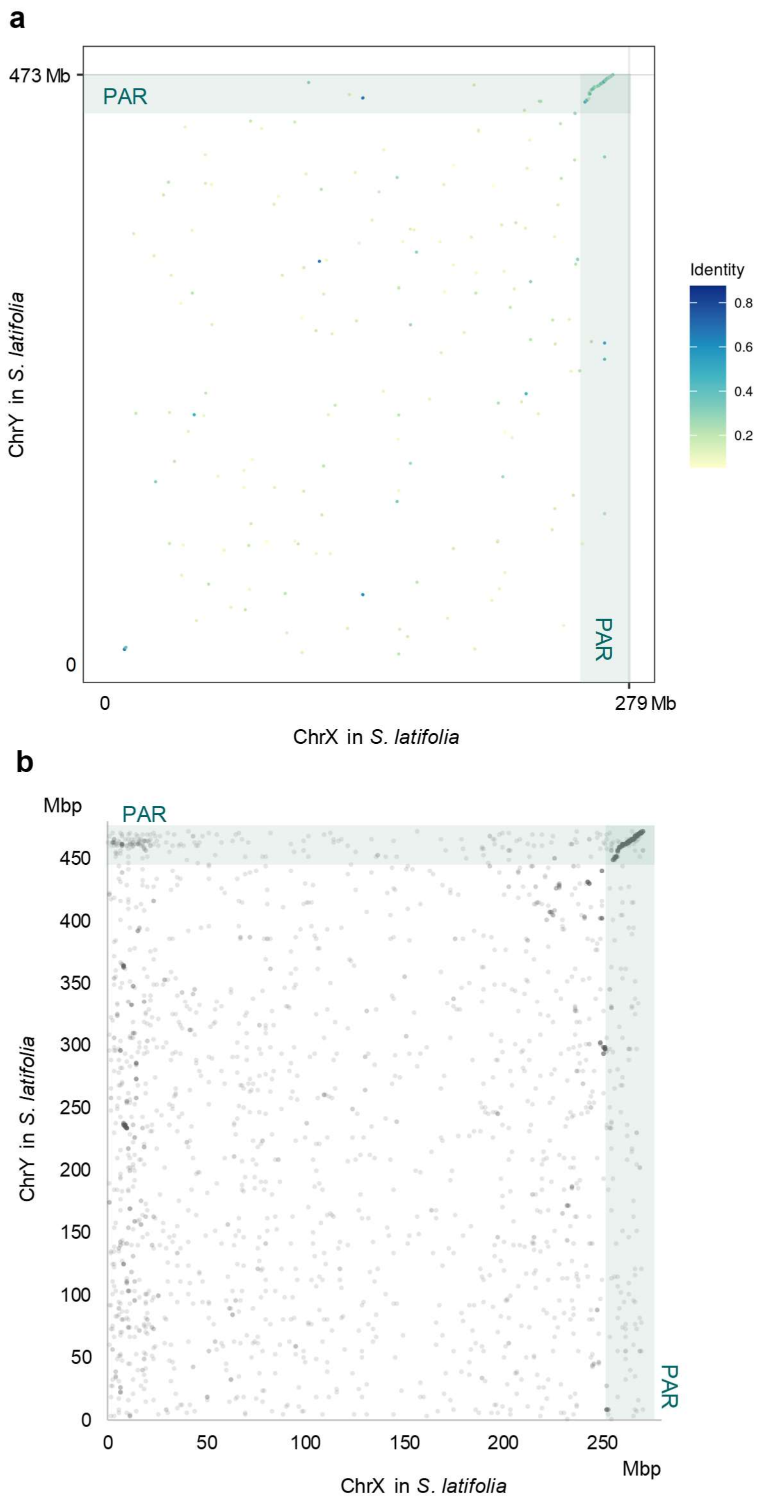


**Figure S11 Sequence-based synteny analysis between the *S. latifolia* X and Y chromosome assemblies**

**a**, Nucleotide sequence-based synteny analysis between X- and Y chromosomes, with MiniMap2. **b**, Relationships of allelic gene positions in X- and Y chromosomes. In both analyses, except the PAR, no clear synteny was conserved between X- and Y chromosomes.

**Figure S12**

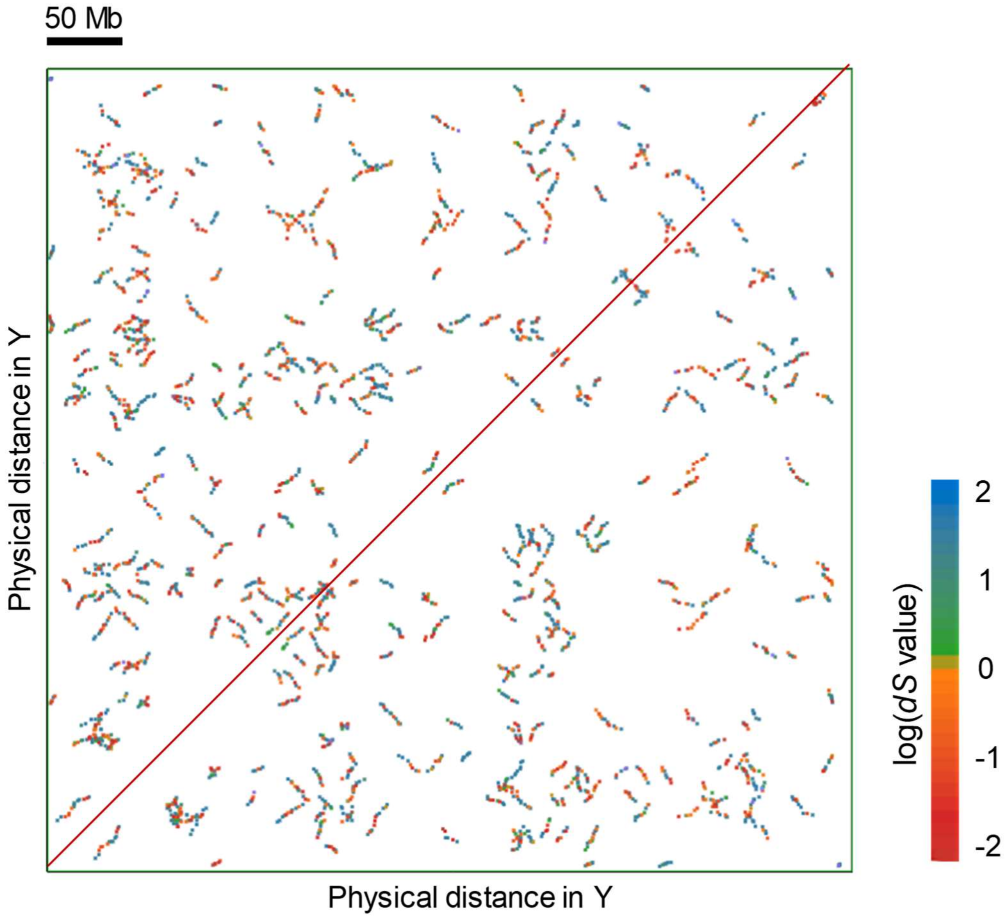


**Figure S12 Gene-order-based self synteny analysis of the *S. latifolia* Y chromosome.** Gene-order based self-synteny analysis was conducted with CoGe (40, 41). The *dS* values of the syntenic blocks were indicated with heatmaps. Y chromosome is constituted of frequent fragmental duplication blocks. Pairwise *dS* values were indicated by the colours shown in the key at the right.

**Figure S13**

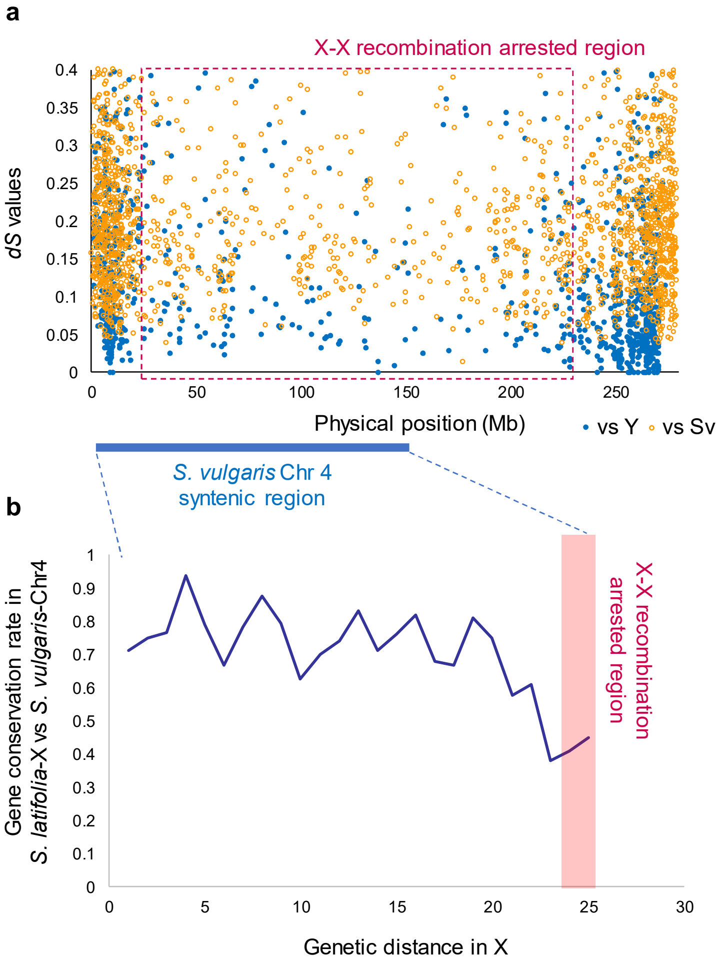


**Figure S13 Distribution of the *dS* values in the physical X chromosome map, and the gene conservation rate in *S. latifolia*-X and *S. vulgaris*-Chr4.**

**a**, Distribution of the *dS* values between X- and Y chromosomal alleles (blue filled circles) and between *S. latifolia* X-chromosomal genes and their orthologs in *S. vulgaris* (orange open circles), in the physical map of the X-chromosome. The approx. 25 - 225 Mb is the recombination arrested region, exhibiting low gene density. **b**, Transition of gene conservation rate in the syntenic region between *S. latifolia* X-chromosome and *S. vulgaris*-Chr4 (see Figure 4 for the details), in the genetic map.

**Figure S14**

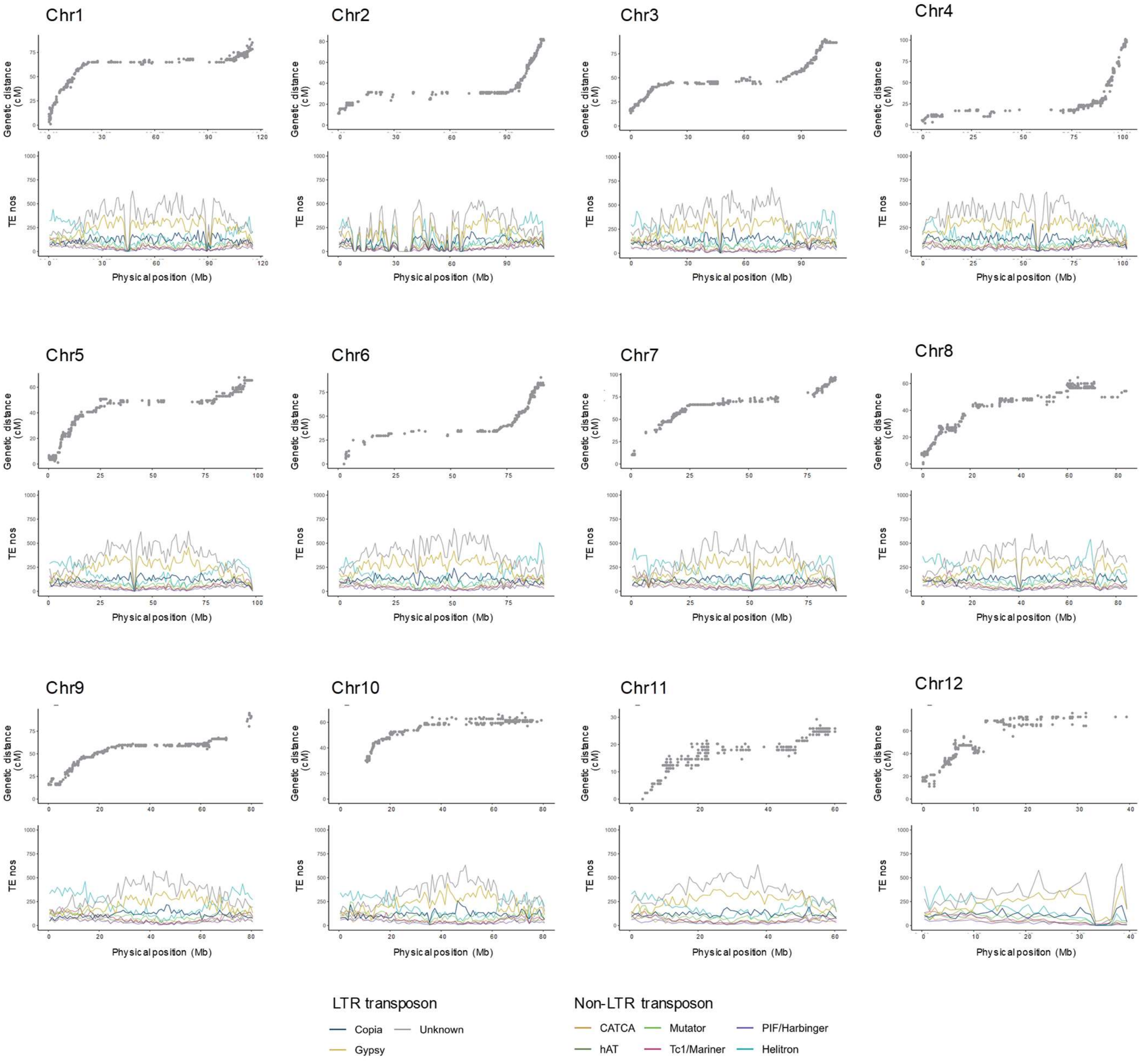


**Figure S14 Genetic recombination rates and distribution of transposons in *S. vulgaris***

Genetic distances according to the physical positions (upper row) and TE distributions (bottom row) were given for 12 chromosomes. For TEs, nine classes were separately provided per 1Mb bin. Consistent with *S. latifolia* (fig. S5), the long recombination arrested regions in each chromosome were highly enriched with LTR-type TEs, especially *Gypsy* and *unknown* classes.

**Figure S15**

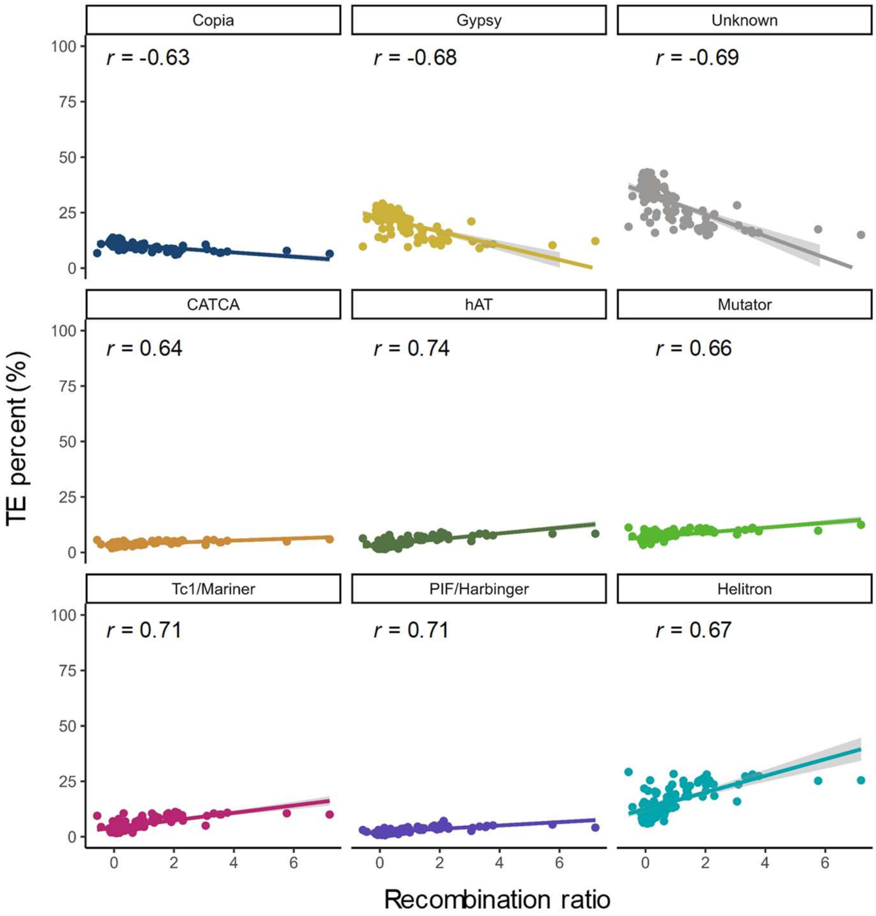


**Figure S15 Correlations between densities of different TE classes and recombination rates in *S. vulgaris***

The Pearson's moment correlation test was performed between the recombination rate (per 10Mb) and each TE class accumulation. Consistent with the trends observed in fig. S14 and in *S. latifolia* (fig. S6), the recombination rates were negatively correlated to LTR-type TEs (or *Copia*, *Gypsy*, and *unknown* classes).

**Figure S16**

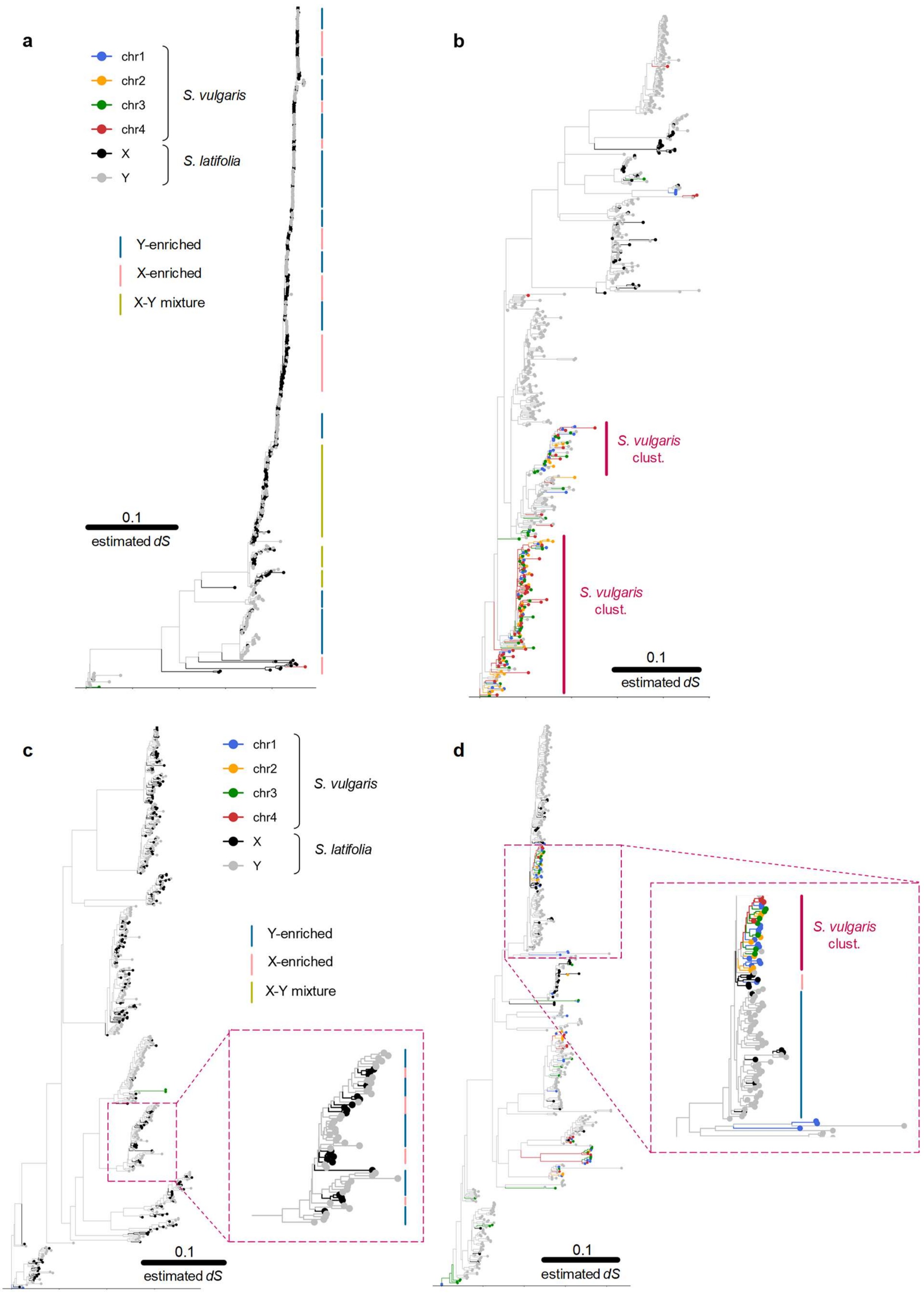
a
chr1
chr2
chr3
chr4
S. vulgaris
X
Y
S. latifolia
Y-enriched
X-enriched
X-Y mixture
0.1
estimated dS
b
S. vulgaris clust.
S. vulgaris clust.
0.1
estimated dS
c
chr1
chr2
chr3
chr4
S. vulgaris
X
Y
S. latifolia
Y-enriched
X-enriched
X-Y mixture
0.1
estimated dS
d
S. vulgaris clust.
0.1
estimated dS

**Figure S16 Representative phylogenies of *Gypsy*- and *unknown*-class LTR-TEs in the sex chromosome assemblies**

Phylogenetic analyses of *Gypsy*-class (**a** and **b**) and *unknown*-class (**c** and **d**) LTR clusters from Fig. 4e-f. **a** and **c**, Clusters enriched with both X and Y components. Although the X and Y components might look randomly located in some clusters, they often underwent recent chromosome-specific duplications in sub-clusters, as given in a dotted box in panel **c**. **b** and **d**, Clusters including both *S. latifolia* and *S. vulgaris* components. Even in clusters including these two species, most components were duplicated in lineage-specific manners, very recently (estimated *dS* << 0.1), as given in a dotted box in panel **d**.

**Figure S17**

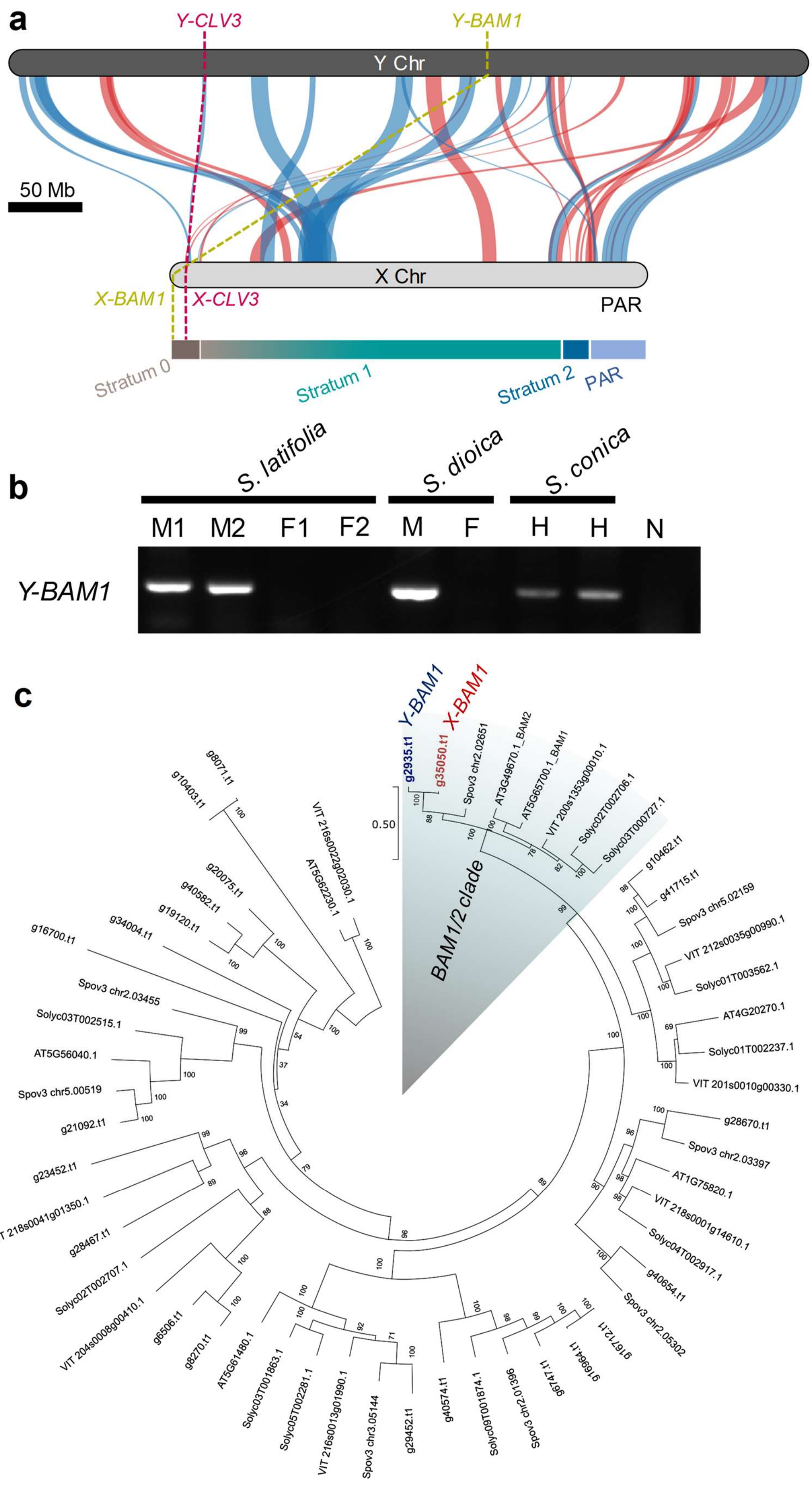
a
Y-CLV3
Y-BAM1
Y Chr
50 Mb
X Chr
X-BAM1
X-CLV3
PAR
Stratum 0
Stratum 1
Stratum 2
PAR
b
S. latifolia
S. dioica
S. conica
M1
M2
F1
F2
M
F
H
H
N
Y-BAM1
c
Y-BAM1
X-BAM1
BAM1/2 clade
0.50

**Figure S17 Characterization of *SlBAM1* gene located on sex chromosomes**

**a**, Physical positions of *Y-BAM1* and *X-BAM1* in Y- and X-chromosomes, respectively. *X-BAM1* is located in the left-hand edge of the oldest stratum (Stratum 0). **b**, Y-BAM1 male-specific sequences were detected in both *S. latifolia* and *S. dioica* (M: male, F: female, M1/F1: in a US line, M2/F2: in a German line), and the gene was also found in hermaphrodite *S. conica* (H: hermaphrodite). We tested with each 5 male and female individuals for independent lines of *S. latifolia* and *S. dioica*. **c**, Phylogenetic relationship of the X/Y-BAM1 with BAM1/2-like genes in angiosperm. X/Y-BAM1 sequences are nested within the monophyletic BAM1/2 family (bootstrap for clade divergence = 100/100). For the OTU prefixes, AT: Arabidopsis, VIT: *Vitis vinifera*, Solyc: *Solanum lycopersicum*, Spov3: *Spinacia oleracea*, g: *Silene latifolia*.

**Figure S18**

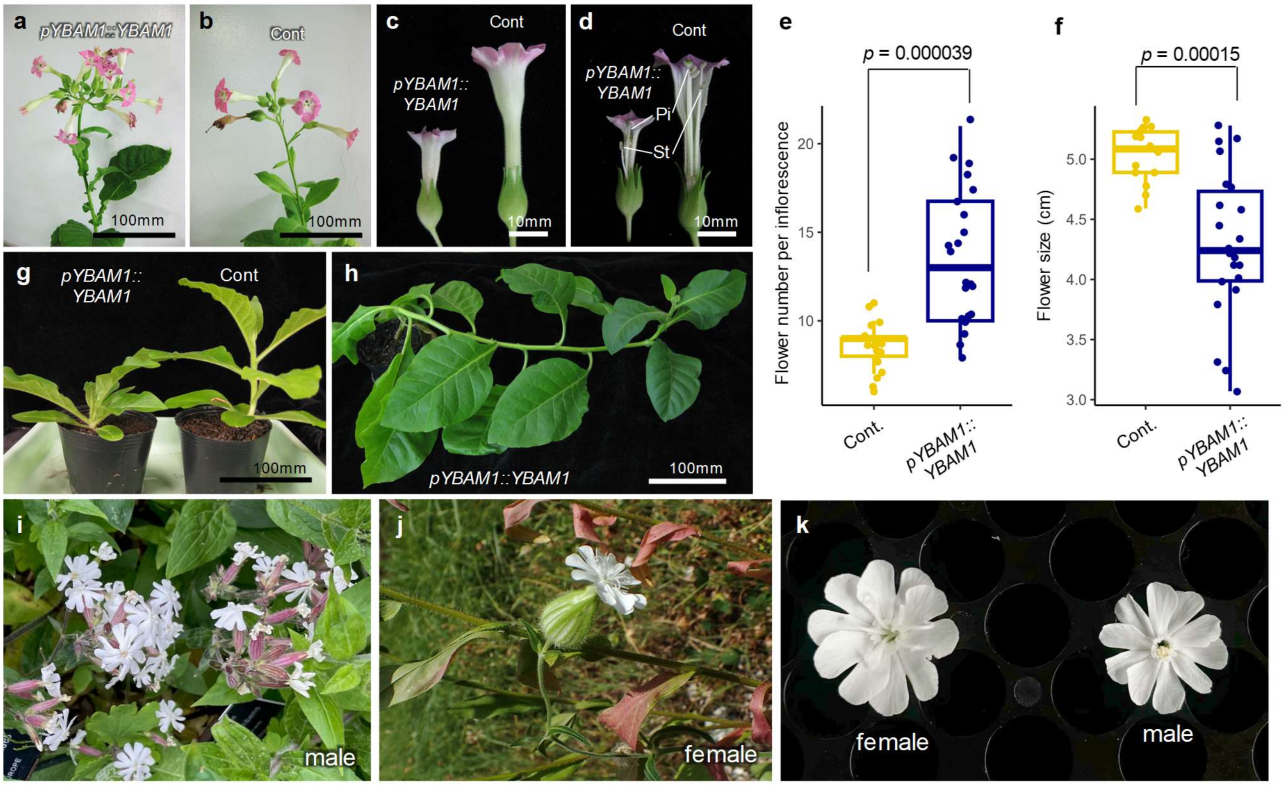


**Figure S18 Functional analysis of *Y-BAM1*, potentially contributing to the sexual dimorphism in *Silene latifolia***

**a-b**, Inflorescence architectures in *Y-BAM1*-introduced (*pYBAM1::YBAM1*) (**a**) and control (Cont) (**b**) *Nicotiana tabacum*. *Y-BAM1* expression was under the control of its native promoter. **c-d**, flower architectures in *Y-BAM1*-introduced and control *Nicotiana tabacum*. Outer appearance (**c**) and dissected (**d**) flowers. **e-f**, Box plots for flower numbers in the first inflorescence (**e**) and flower sizes in the first five flowers (**f**). The flower numbers were substantially increased in the *pYBAM1::YBAM1* lines, in comparison to the control lines ($p$ = 0.000039 in two-sided Student's $t$-test). On the other hand, flower size in the *pYBAM1::YBAM1* lines was smaller than the control lines ($p$ = 0.00015 in two-sided Student's $t$-test). **g**, The *pYBAM1::YBAM1* lines often exhibited shorter nodes than the controls. **h**, A few of the *pYBAM1::YBAM1* lines showed continuous vegetative growth (or prolonged juvenile phase) and substantial delay in flowering. Panel **h** showed a 5-month age *pYBAM1::YBAM1* line, without any flower buds. **i-j**, Representative inflorescence architectures in male (**i**) and female (**j**) *S. latifolia*. **k**, Representative male and female flowers in *S. latifolia*. Their male phenotypes, more flowers per inflorescence and smaller flower size, are consistent with the effects of *Y-BAM1*.

**Figure S19**

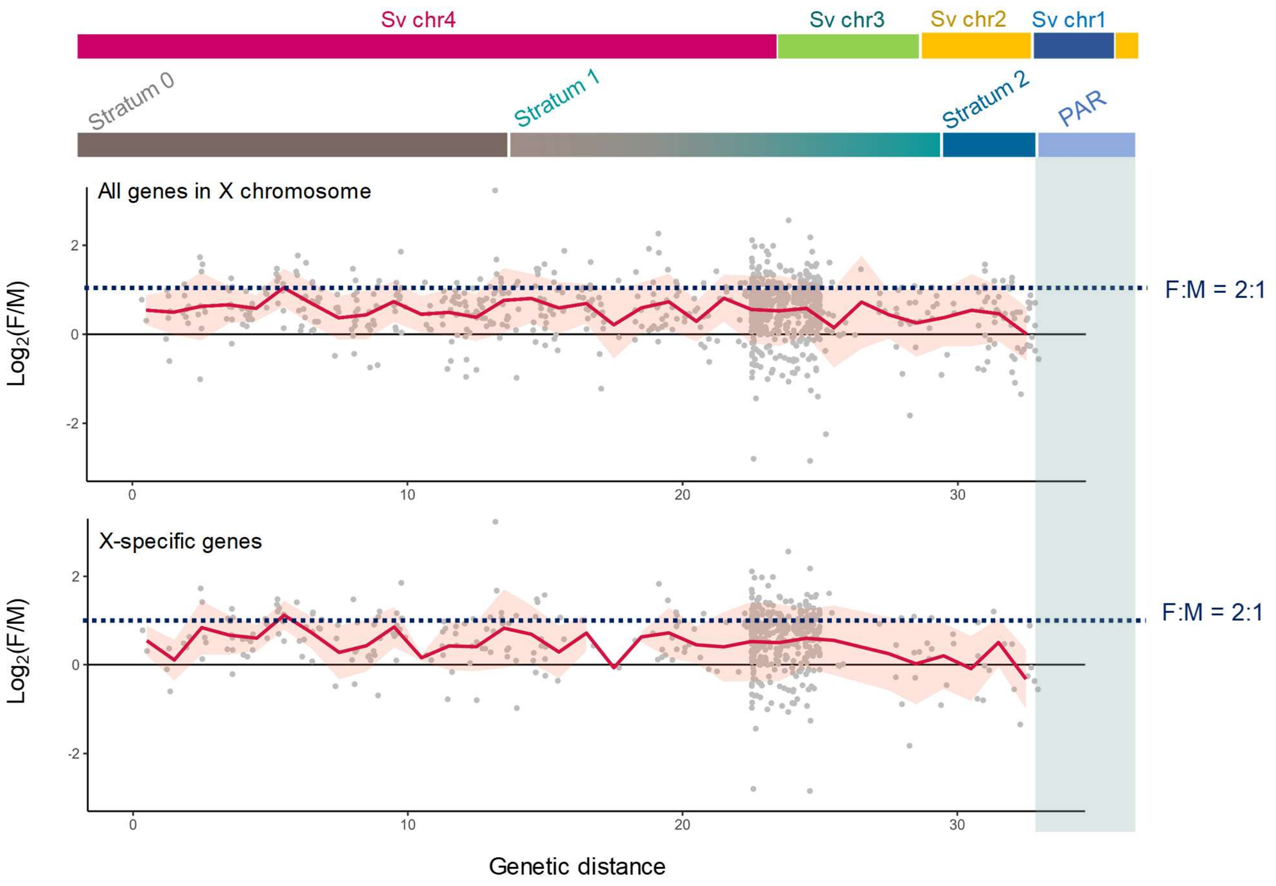


**Figure S19 Test for X chromosome dosage compensation in *S. latifolia***

Gene expression levels in young leaves ($5^{th}$ leaves under identical growth condition) were compared in male and female offspring of the same parents, with *N* = 4 biological replicates. The two plots showing the results are for two analyses, one using all X-chromosomal genes, and the other using X-specific genes. Each dot corresponds to one gene, and the genes' positions are shown on the x axis, using the genetic map positions in female meiosis. As in the main text Figure 3j, many genes in the pericentomeric region are within a small genetic map distance just before 25 cM. The red lines indicate the mean values in 1cM bins, and the pink ribbons show one standard deviation. The values for most genes were within the $\log_2$(female/male) range 0-1, suggesting that some X-linked genes exhibit dosage compensation. There was no relationship between this measure of dosage compensation and the distance to the PAR, though the results for X-specific genes tend to be low for the youngest stratum, 2).

#### Figure S20

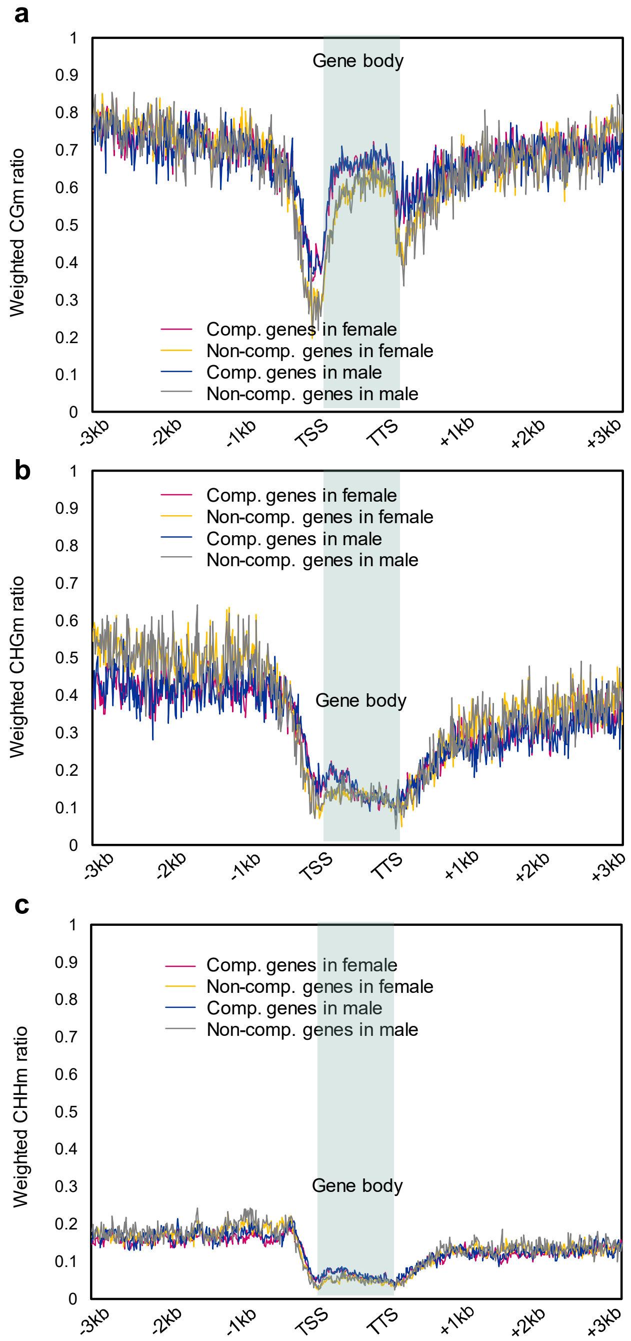


**Figure S20 DNA methylation profiles in the genes exhibiting X chromosome dosage compensation**

DNA methylation levels in the three contexts, CG (**a**), CHG (**b**), and CHG (**c**). The panels show results in in females and males, for sequences surrounding two categories of X-linked genes: dosage-compensated (Comp.), defined as showing female/male expression < 1.5, non-dosage-compensated (No-comp.) The CG and CHG methylation patterns differ slightly between Comp. and Non-comp genes, but not between the sexes. Hence, DNA methylation, suppressing X-linked gene expression in females, cannot be the mechanism of dosage compensation in this plant.
